# A molecular network of conserved factors keeps ribosomes dormant in the egg

**DOI:** 10.1101/2021.11.03.467131

**Authors:** Friederike Leesch, Laura Lorenzo-Orts, Carina Pribitzer, Irina Grishkovskaya, Manuel Matzinger, Elisabeth Roitinger, Katarina Belačić, Susanne Kandolf, Tzi-Yang Lin, Karl Mechtler, Anton Meinhart, David Haselbach, Andrea Pauli

## Abstract

Ribosomes are produced in large quantities during oogenesis and stored in the egg. However, the egg and early embryo are translationally repressed. Using mass-spectrometry and cryo-EM analyses of ribosomes isolated from zebrafish and *Xenopus* eggs and embryos, we provide molecular evidence that ribosomes transition from a dormant to an active state during the first hours of embryogenesis. Dormant ribosomes are associated with four conserved factors that form two modules and occupy functionally important sites of the ribosome: a Habp4-eEF2 module that stabilizes ribosome levels and a Dap1b/Dapl1-eIF5a module that represses translation. Dap1b/Dapl1 is a newly discovered translational inhibitor that stably inserts into the polypeptide exit tunnel. Thus, a developmentally programmed, conserved ribosome state plays a key role in ribosome storage and translational repression in the egg.

## Introduction

Ribosomes are amongst the most abundant macromolecular complexes stored in the quiescent egg^1,2^. These maternally provided ribosomes (in short: maternal ribosomes) are essential for progression through embryogenesis as they are required for the translation of maternal and zygotic transcripts^3–6^ (Fig. 1a). Although the overall number of ribosomes per embryo remains constant throughout early embryogenesis, prior studies from multiple organisms indicate that translational activity increases in the embryo, suggesting that translation is repressed in the mature egg^7–12^. Several mechanisms, including shortening of mRNA polyadenine tails^13^ and interference with the formation of the translational initiation factor eIF4F^14,15^, have been implicated in dampening translation in the egg and early embryo, while the ribosome itself has not been considered to contribute to this regulation.

**Fig. 1.**
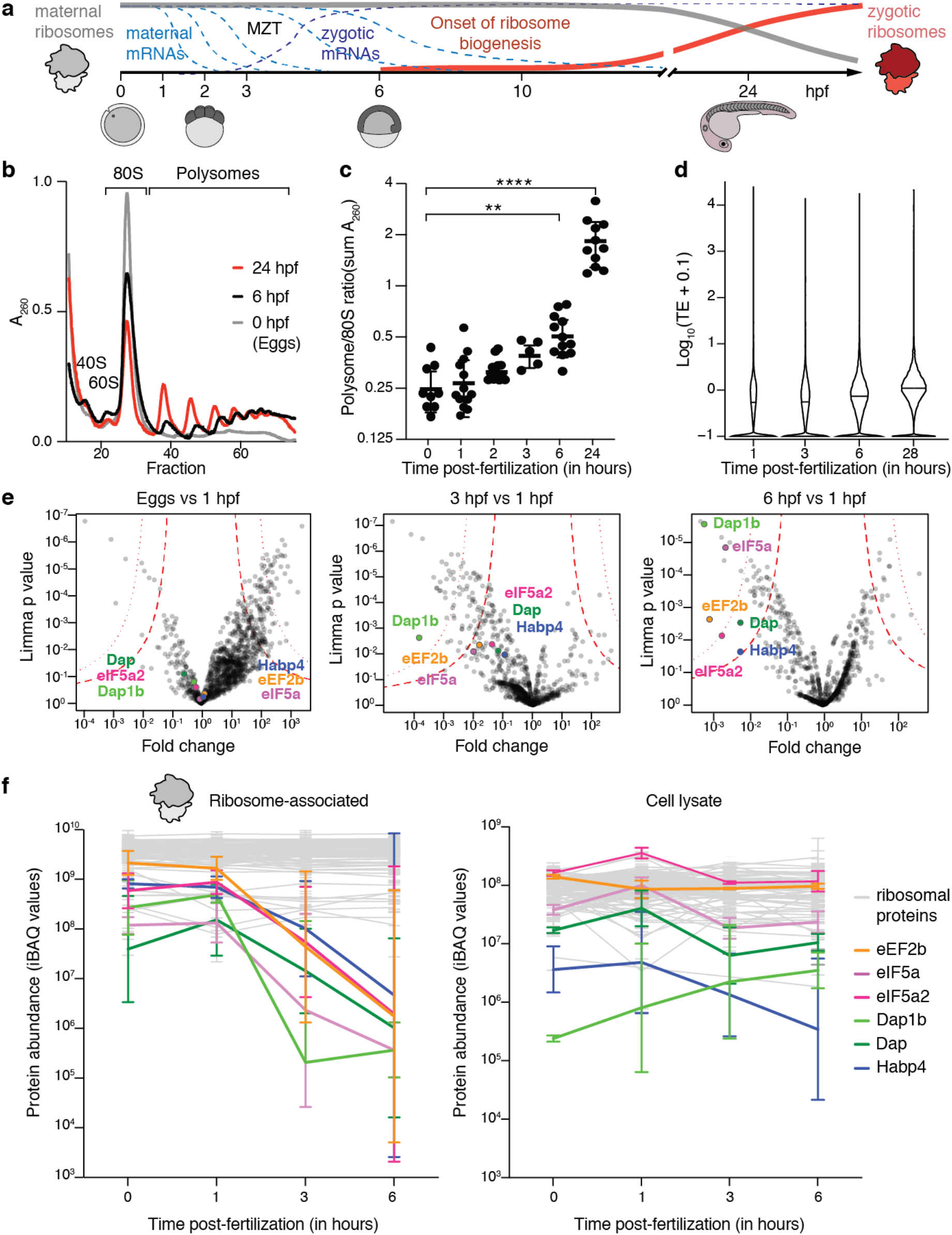
Increase of translation during zebrafish embryogenesis correlates negatively with the presence of ribosome-bound factors. **a**, Schematic of the maternal-to-zygotic transition (MZT) in zebrafish, in which the clearance of maternal mRNAs (light blue dashed line) is coordinated with the activation of zygotic mRNA transcription (dark blue dashed line) during the first hours of development. The absolute time (in hours post-fertilization (hpf)) is indicated below the graph, along with schematics of embryos. The replacement of maternal ribosomes (grey) by zygotic ribosomes (red) is delayed compared to the MZT and occurs over the course of several days. **b**, Representative polysome profiles of zebrafish eggs, 6 hpf embryos and 24 hpf larvae. A260, absorbance value at 260 nm. **c**, Quantification of polysome-to-monosome ratios at different stages of development (0 h: n=8; 1 h: n=12; 2 h: n=5, 3 h: n=6; 6 h: n=12; 24 h: n=11) (**: *p-value* = 0.0013, ***: *p-value* > 0.0001). **d**, Violin plots showing the distribution of the median translation efficiency (TE), calculated as the ratio between ribosome protected fragments and total RNA counts, at different developmental time points. **e**, Volcano plots based on mass spectrometry analysis showing fold enrichments of zebrafish proteins in the ribosome fraction of 1 hpf zebrafish embryos compared to eggs (left), 3 hpf embryos (middle) and 6 hpf embryos (right) (n=3). Permutation-based false discovery rates (FDRs) are displayed as dotted (FDR < 0.01) and dashed (FDR < 0.05) lines. Ribosome-associated factors that are stable in eggs and 1 hpf stages but depleted from ribosomes during the first 6 hours of embryogenesis are highlighted in color. **f**, Abundance changes of a subset of factors relative to core ribosomal proteins in the ribosome-associated proteome (left) and total cell lysate (right) during the egg-to-embryo transition (n=3). Protein abundances are reported as iBAQ values (sum of all peptide intensities divided by the number of observable peptides for a given protein). Core ribosomal proteins are plotted in light grey (76/81: left; 74/81: right); ribosome-associated factors that are released from the ribosome are highlighted in color.

Translationally inactive ribosomes have been observed in pro- and eukaryotes in specific cellular contexts, e.g. immunity, viral infection, and nutrient deprivation, and are associated with factors such as Bac7^16^, Nsp1^17^, and Stm1^18^, respectively. Studies from the 1970s with sea urchin eggs indicated the presence of inhibitory proteins which were thought to be associated with maternal ribosomes^19–21^, yet the relevance and molecular identity of these factors remained unclear. How ribosomes are stored in an inactive state for extended amounts of time in the mature egg, and whether regulation of the ribosome itself contributes to translational repression in the egg and the subsequent increase in translational activity during embryogenesis remains thus unknown.

## Results

To investigate the timing of translational activation during the egg-to-embryo transition in vertebrates, we examined the translational status of zebrafish embryos at different stages of development (Fig. 1a). Specifically, we used polysome gradients to uncover the fraction of ribosomes present as individual subunits (40S and 60S), monosomes (80S) and polysomes. In the absence of triggers of ribosome stalling, an increase in the polysome fraction generally indicates increased active translation^22^. We found that ribosomes in the egg and in the 1-hour post fertilization (hpf) embryo were almost exclusively present as monosomes, whereas the polysome fraction started to increase from 3-6 hpf onwards (Fig. 1b, c). As an orthogonal approach, we calculated translational efficiency (TE) values based on ribosome-protected mRNA fragments over the course of zebrafish embryogenesis, using published ribosome profiling and RNA-Seq data sets^23,24^. We observed that translational efficiency increases over the course of embryogenesis, with a clear increase at 6 hpf (Fig. 1d). Together, these data suggest that, similarly to other vertebrates^7–12^, most ribosomes present in the zebrafish egg and early embryo were not engaged in active translation. Given that zygotic ribosomes do not accumulate until after 6 hpf (Fig. 1a)^3,4,25^, we hypothesize that maternal ribosomes transition from a “dormant” to an “active” state, leading to increased translational activity during the first hours of embryogenesis in zebrafish.

To uncover factors that regulate maternal ribosome activity during the first hours after fertilization, we purified bulk ribosomes (monosomes and polysomes) from zebrafish eggs and developing embryos for mass-spectrometry (MS) analysis. While the relative abundances of core ribosomal proteins were similar at all time points, we identified four evolutionarily conserved factors, namely eIF5a/eIF5a2, eEF2b, Habp4 and the paralogs Dap1b/Dap, that were specifically associated with ribosomes in zebrafish eggs and 1 hpf embryos, but depleted from ribosomes in 3 hpf and 6 hpf embryos (Fig. 1e, f). The levels of these ribosome-associated factors remained relatively stable in total embryo lysates over the first 6 hours of development (Fig. 1f), suggesting that they were released from the ribosome but not immediately degraded. Importantly, we found that the same set of factors were enriched in ribosomes purified from *Xenopus* eggs versus 24 hpf embryos (a developmental stage equivalent to zebrafish 6 hpf embryos) (Extended Data Fig. 1a). Thus, the association of these four factors with ribosomes in the mature egg is conserved in zebrafish and *Xenopus* and correlates with suppressed translation.

To elucidate how these factors establish a translationally repressed ribosome state, we determined the structures of maternal ribosomes isolated from 1 hpf zebrafish embryos (Fig. 2a) and *Xenopus* eggs (Fig. 2b) by electron cryo-microscopy (cryo-EM) at 3.2 and 2.8 Å average resolution, respectively (Extended Data Table 1, Extended Data Figs. 2, 3). We were able to assign densities for all four factors in ribosomes from both zebrafish and *Xenopus* (Fig. 2c, Extended Data Fig. 1b-e,). In parallel, we performed cryo-EM analysis of ribosomes isolated from 6 hpf zebrafish embryos (2.6 Å average resolution; Extended Data Figs. 1f, 4, Extended Data Table 1). In agreement with the MS data, maternal ribosomes from 6 hpf zebrafish embryos showed low density levels for eEF2b and no densities for eIF5a/eIF5a2, Habp4 and Dap1b/Dap (Extended Data Fig. 1f, Extended Data Table 2).

**Fig. 2.**
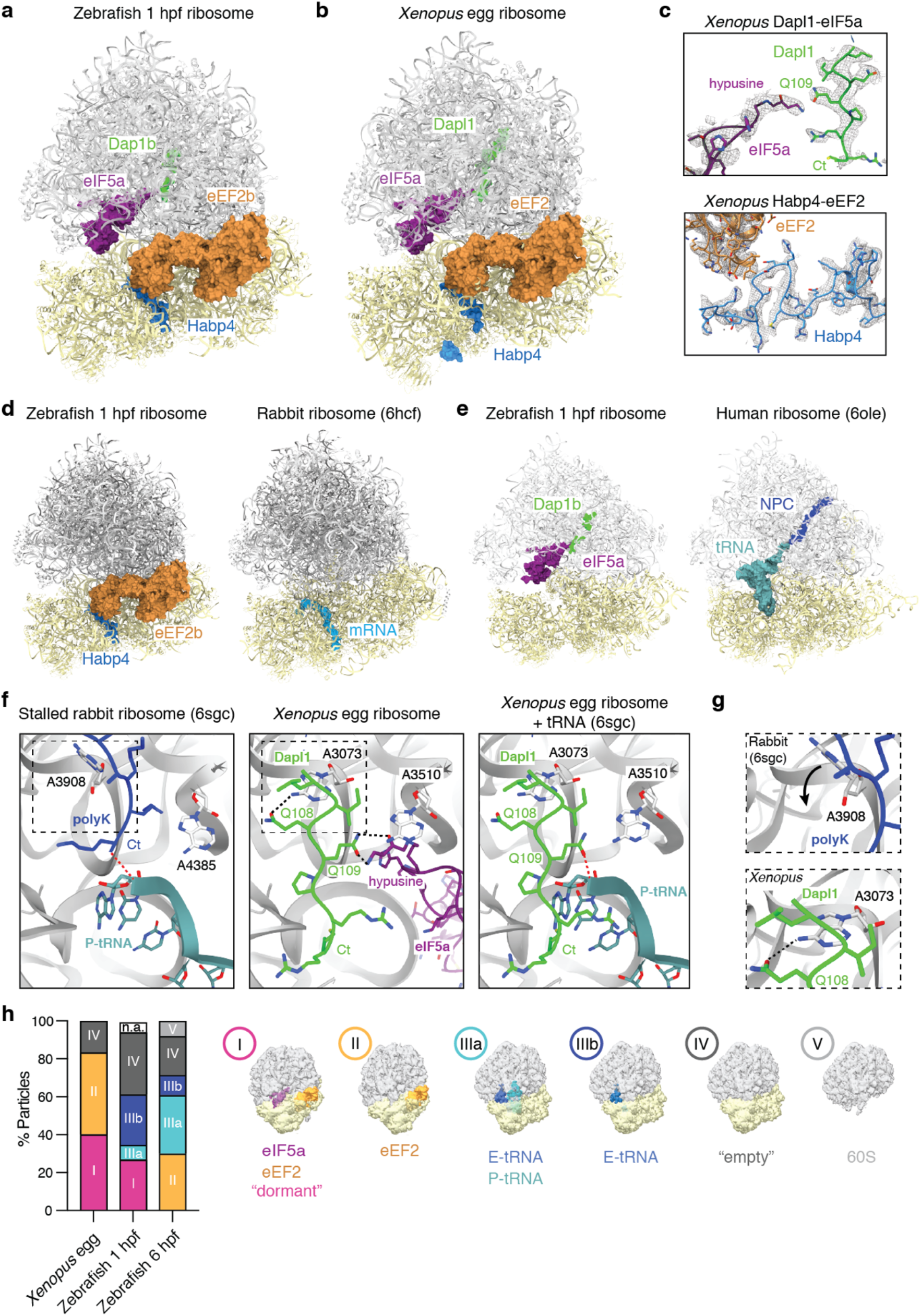
A conserved set of factors blocks functionally important sites of the egg-state ribosome. **a-b**, Overview of the dormant ribosome structure from 1 hpf zebrafish (a) and *Xenopus* egg (b). Core ribosomal proteins and rRNA molecules are shown in gray (large subunit) and yellow (small subunit). Ribosome-associated factors are shown as surface representations; eEF2b (in orange) and eIF5a (in dark magenta) correspond to PDB-6MTE^31^ and PDB-5DAT, and were aligned to ribosomes using the command *mmaker* in ChimeraX (root-mean-square deviations (RMSDs) for alignment to 1 hpf zebrafish ribosome: 0.454 Å and 0.643 Å, respectively; RMSDs for alignment to *Xenopus* egg ribosome: 0.489 Å and 0.642 Å, respectively). **c**, Density maps (in mesh) of the two identified modules of the *Xenopus* dormant ribosome: Dapl1-eIF5a (top) and Habp4-eEF2 (bottom). The distance between the carboxyl group of Q109 of Dapl1 and the amine group of hypusine-51 of eIF5a is 2.9 Å (2.5 Å in the zebrafish 1 hpf ribosome between Q105 of Dap1b and hypusine-51). Ct, C-terminus. **d**, Habp4 (in blue) and eEF2b (in orange, from 6MTE^31^) are shown as surface representations in the dormant ribosome from 1 hpf zebrafish (left). The structure of the rabbit ribosome stalled on an mRNA (depicted as surface representation in light blue; 6HCF^42^ is shown on the right for comparison. Both structures were superimposed using the command *mmaker* in ChimeraX (RMSD: 0.957 Å). **e**, Clipping of the dormant ribosome from 1 hpf zebrafish shows Dap1b (in green) bound within the polypeptide exit tunnel (PET) (left). The structure of a stalled human ribosome, 6OLE^43^, containing a nascent polypeptide chain (NPC, in blue) and a P-tRNA (in turquoise) is shown on the right for comparison. Dap1b, eIF5a (in dark magenta; from 5DAT), the NPC and the P-tRNA are depicted as surface representations. Both structures were superimposed using the command *mmaker* in ChimeraX (RMSD: 0.689 Å). **f**, Comparison of the peptidyl-transferase center (PTC) of a rabbit 80S ribosome stalled with a poly-Lysine NPC (in blue) (6SGC^39^, left) and of the *Xenopus* egg ribosome. The large subunits of both models were superimposed using the command *mmaker* in ChimeraX (RMSD: 0.297Å). The P-tRNA (in turquoise) (6SGC^39^) is shown superimposed onto the *Xenopus* egg ribosome for comparison on the right (eIF5A is hidden). Critical amino acids and 28S rRNA nucleotides are labelled; interactions are depicted with dashed lines. Boxed areas (dashed boxes) are shown at higher magnification in g. **g**, Magnified view of the boxed areas in F. A3073 of *Xenopus* 28S rRNA (bottom, equivalent to A3908 of rabbit 28S on top) displays a different conformation when interacting with Gln108 of Dapl1. **h**, Distribution of ribosomal particles among the classes obtained after the analysis of particle heterogeneity in *Xenopus* egg ribosomes and zebrafish 1 hpf and 6 hpf ribosomes with cryoDRGN^44^. Representative filtered density maps of the six major classes are shown on the right (see Extended Data Table 2; n.a refers to “non-assigned” particles).

eIF5a promotes translational elongation and termination, particularly upon ribosome stalling at specific amino acid sequence contexts^26,27^. We found that eIF5a/eIF5a2 binds between the exit (E) and peptidyl (P) site of the ribosome, as previously reported^28^. eEF2 mediates ribosomal translocation^29,30^ and transiently interacts with the tRNA-mRNA complex at the aminoacyl (A) site of the ribosome. Proteins of the Habp4 family (PFAM family PF04774), but not Habp4 itself, have been reported to stabilize the eEF2 interaction at the A site in inactive ribosomes^31,32^. Indeed, we found Habp4 bound to the mRNA entry channel of zebrafish 1 hpf and *Xenopus* egg ribosomes at the same position as previously reported for its mammalian homolog SERBP1^31,33^, sequestering eEF2 at the A site and blocking the interaction sites of the tRNA-mRNA complex on the ribosome (Fig. 2d; Extended Data Fig. 1b, c).

Death Associated Protein 1b (Dap1b)/Dapl1 (in *Xenopus* Dapl1.L/Dapl1.S; in mammals also known as EEDA (early epithelial differentiation-associated)^34,35^) and Dap are highly conserved proteins across animals (Extended Data Fig. 5) but have not been reported to associate with the ribosome. In mammals, Dapl1 has been linked to epithelial differentiation and proliferation^34– 36^, and Dap has been described as a positive regulator of programmed cell death^37^ and a negative regulator of autophagy^38^. However, the molecular mechanisms of action for both factors are still unknown. Strikingly, we found density in the polypeptide exit tunnel (PET) of zebrafish 1 hpf and *Xenopus* egg ribosomes that we could attribute to the conserved C-terminus of Dap1b/Dapl1 or Dap (Fig. 2e, Extended Data Figs. 1d, e, 5). Dap1b was ∼10-fold more abundant than Dap in ribosomes from 1 hpf zebrafish embryos according to our MS analysis (Fig. 1f), and only Dapl1 was identified by MS in ribosomes from *Xenopus* eggs (Extended Data Fig. 1a). Thus, we modelled the C-terminus of zebrafish Dap1b and *Xenopus* Dapl1, which we found to extend by five amino acids beyond the position that is occupied by the C-terminal amino acid residue of a nascent polypeptide chain (Fig. 2c, f). Intriguingly, an invariant Glutamine in Dap1b/Dapl1 (Gln109 in *Xenopus*, Gln105 in zebrafish, Extended Data Fig. 5) occupies the same position as the C-terminal residue of a nascent peptide chain^39^ and forms a hydrogen bond with the hypusine residue of eIF5a (Fig. 2f). Of note, hypusination of eIF5a’s Lysine residue is essential for eIF5a function^40,41^. Moreover, we observe additional hydrogen-bond interactions between the immediately adjacent Glutamine in Dap1b/Dapl1 (Gln108 in *Xenopus*, Gln104 in zebrafish, Extended Data Fig. 5) with the 28S rRNA (Fig. 2f, g). Interestingly, the interaction of *Xenopus* Gln108 (Gln104 in zebrafish) with adenine 3073 (A3168 in zebrafish) restricts this adenine to a conformation distinct from previously reported ribosomal structures (Fig. 2g). Together, our structural data suggests that Dap1b/Dapl1’s specific interactions with eIF5a and the 28S rRNA contribute to Dap1b/Dapl1’s binding to the PET of a fully assembled eukaryotic 80S ribosome, thus making them unique among all ribosome-associated factors characterized so far.

To determine whether the four factors are bound to the same ribosomal particles, we performed a 3D variability analysis after dimensional reduction of the datasets using cryoDRGN^44^. Each dataset’s latent representation showed distinct clusters when visualized in UMAPs (Extended Data Fig. 1g, h). A detailed analysis of the particles in these clusters revealed six major classes based on the presence or absence of eEF2, eIF5a, and tRNAs (Extended Data Table 2). In contrast to previous reports suggesting mutually exclusive binding of eIF5a and eEF2 to the ribosome^26,31,45^, our analysis revealed that 40.2% and 26.9% of ribosomes in *Xenopus* and in 1 hpf zebrafish, respectively, were simultaneously bound by eIF5a and eEF2 (Fig. 2h, class I particles). Due to the dimensional reduction of the datasets, cryoDRGN-derived maps were not of sufficient resolution to confidently assess the presence of Dap1b/Dap and Habp4 in any of the clusters. However, given the data from the literature on Habp4 homologs binding to eEF2-containing ribosomes^31–33^ and our data showing that eIF5a and Dap1b interact with each other (Fig. 2c, f), our results are consistent with Dap1b and Habp4 being bound to class I (eIF5a- and eEF2-containing) ribosomes (Fig. 2h). In support of this conclusion, class I particles were not observed in the translationally more active 6 hpf zebrafish embryos (Fig. 2h, Extended Data Table 2, Extended Data Fig. 1h). In contrast to class I particles, which were exclusively identified in the translationally repressed zebrafish and *Xenopus* egg states and will thus in the following be referred to as “dormant” state, other ribosomal particle classes were identified to varying levels in all data sets, including in 6 hpf zebrafish embryos (Fig. 2h, Extended Data Fig. 1h). Other particle classes include class II particles that contained only eEF2, class III particles that contained tRNAs, and “empty” class IV particles that lack eEF2, eIF5a, and tRNAs (Fig. 2h). “Empty” class IV particles may correspond to ribosomes which have lost ribosome-associated factors or tRNAs during the purification since they do not cluster together but occupy different positions in the UMAPs (Extended Data Fig. 1h). Together, our results support a novel, dormant ribosome state present in the mature egg in which four functionally important sites of the ribosome, namely the A site and the E/P site, the mRNA channel and the PET are occupied by evolutionarily conserved factors.

While we could assign Cryo-EM densities for small amino acid stretches within Habp4 and Dap1b/Dap1l and Dap proteins, the majority of the polypeptide chains are not visible in our ribosome structures. To determine the path of these proteins in relation to ribosomal proteins, as well as to confirm that Dap1b/Dapl1 and Dap are novel ribosome-associated factors, we performed crosslinking MS with the primary amine crosslinker disuccinimidyl sulfoxide (DSSO), using ribosomes purified from 1 hpf zebrafish embryos and *Xenopus* eggs (Fig. 3a-d, Extended Data Tables 3-5). We obtained more than 1000 crosslinked peptides for each ribosome sample. 95% (zebrafish) and 90% (*Xenopus*) of all mapped crosslinks had Cα-Cα distances below the expected maximum crosslinking length of 23 Å (Extended Data Fig. 1i), revealing high quality of our crosslinking MS data^46^. We found crosslinks of the N-terminus of Habp4, which was not visible in cryo-EM densities, with small and large subunit ribosomal proteins, including Rpl7a and Rps3a, in dormant ribosomes from both zebrafish and *Xenopus* (Fig. 3a, b). Crosslinking MS analyses of Dap1b/Dapl1 and Dap showed that the N-terminal regions are proximal to the polypeptide exit site, which is consistent with insertion of their C-termini into the PET. In particular, we found crosslinks between a highly conserved N-terminal region of Dapl1/Dap and Rpl31, and between the central region of Dapl1/Dap and Rpl35 (Fig. 3c, d; Extended Data Fig. 5), indicating that both N- and C-terminal regions of Dap1b/Dapl1 and Dap are in close proximity to the ribosome.

**Fig. 3.**
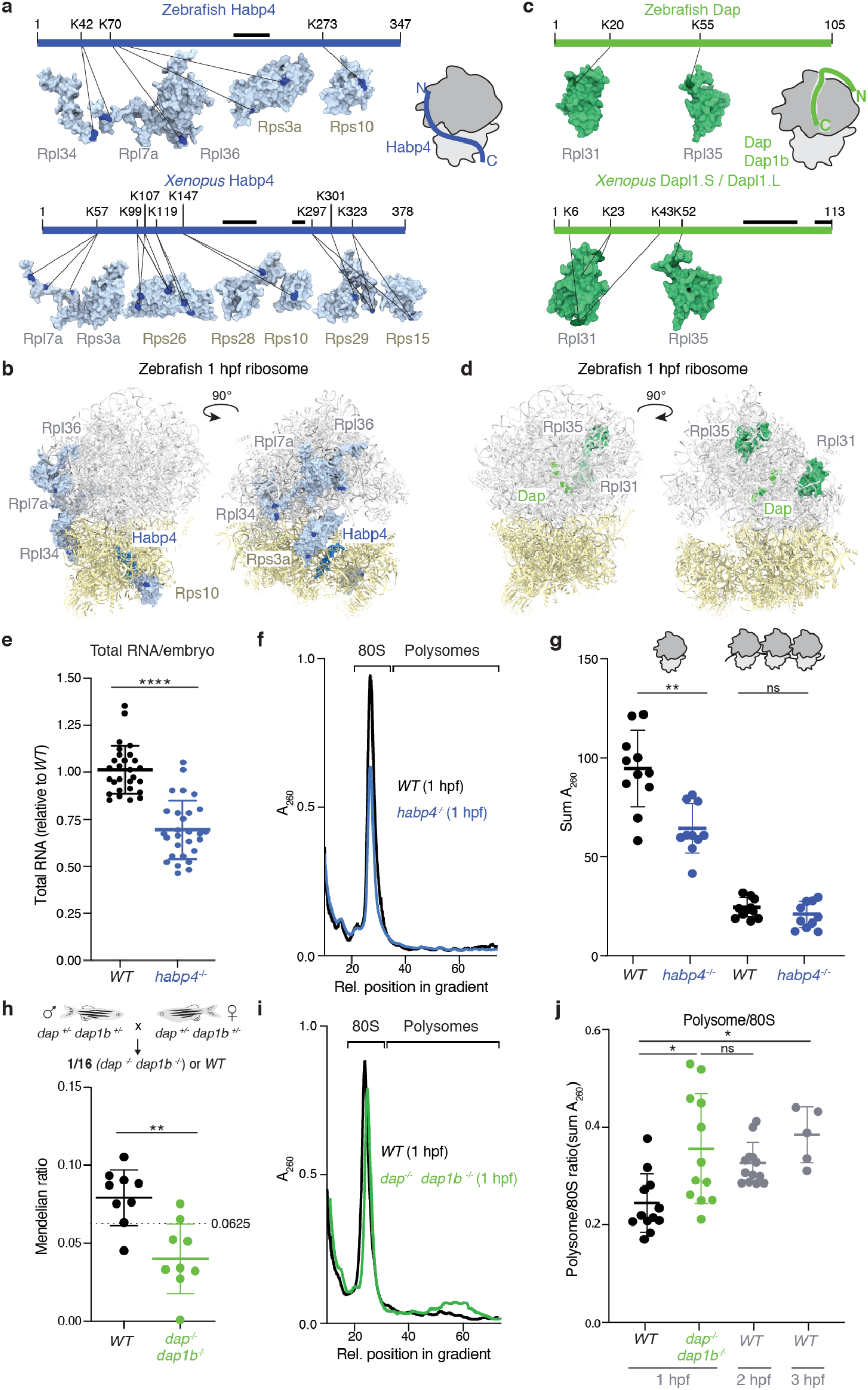
Habp4 binding to the mRNA channel stabilizes monosomes in the egg, while Dap1b/Dap binding to the polypeptide exit tunnel represses polysome formation in zebrafish embryos. **a**, Crosslinking mapping of Habp4 (depicted as a scheme; modeled regions highlighted with a black line) to proteins of the 1 hpf zebrafish (top) and *Xenopus* egg (bottom) ribosomes. Crosslinked proteins are shown as surface representations, with crosslinked residues depicted in dark blue. A cartoon on the top right shows the proposed position of Habp4 (in blue) on the ribosome based on crosslinking and cryo-EM data. **b**, Position of the proteins crosslinked to zebrafish Habp4 (shown as surface representations) in the 1 hpf zebrafish ribosome. **c**, Crosslinking mapping of Dap and Dapl1 (depicted as a scheme; modeled regions highlighted with a black line) to proteins of the 1 hpf zebrafish (top) and *Xenopus* egg (bottom) ribosomes, respectively. Crosslinked proteins are shown as surface representations, with crosslinked residues depicted in dark green. A cartoon on the top right shows the proposed position of Dapl1/Dap (in green) on the ribosome based on crosslinking and cryo-EM data. **d**, Position of the proteins crosslinked to zebrafish Dap (shown as surface representations) in the 1 hpf zebrafish ribosome. **e**, Comparison of the amount of total RNA per embryo, normalized to the average total RNA amount per wild-type embryo, from 1-3 hpf embryos derived from wild-type (*WT*) and *habp4*^*-/-*^ parents (n=28, *p-value <* 0.0001). **f**, Representative polysome profiles from 1 hpf embryos derived from *WT* and *habp4*^*-/-*^ parents. A260, absorbance value at 260 nm. **g**, Quantification of the monosome (left) and polysome (right) peaks of polysome profiles from 1 hpf embryos derived from *WT* and *habp4*^*-/-*^ parents (n=10, *p-value* (monosome comparison) = 0.0056; ns, not significant). **h**, Mendelian ratio of fin-clipped adult fish. Adult double homozygous *dap*^*-/-*^, *dap1b*^-/-^ fish as well as *WT* fish are expected at 6.25% (1/16 fish) from an in-cross of double-heterozygous *dap*^*+/-*^, *dap1b*^*+/-*^ parents (n=9 crosses; min. 84 fish analyzed per cross; total number of fish genotyped: 1029). **i**, Representative polysome profiles from 1 hpf embryos derived from *WT* and *dap*^*-/-*^, *dap1b*^*-/-*^ parents. **j**, Quantification of polysome-to-monosome ratios of polysome profiles from 1 hpf embryos derived from *WT* and *dap*^*-/-*^, *dap1b*^*-/-*^ parents (n=12, *p-value [1 hpf WT vs 1 hpf dap*^*-/-*^, *dap1b*^*-/-*^] = 0.0156; *p-value* [2 hpf *WT* vs 1 hpf *dap*^*-/-*^, *dap1b*^*-/-*^] = ns, not significant; *p-value* [1 hpf *WT* vs 3 hpf *WT*] = 0.0047).

Zebrafish and *Xenopus habp4, dap1b/dapl1* and *dap* mRNAs are highly expressed during oogenesis, whereas *eif5a* and *eef2* transcripts are abundant in all tissues (Extended Data Fig. 6). Moreover, Habp4 family proteins have been linked to ribosome stabilization during nutrient deprivation in yeast (Stm1)^47^ and to non-translating ribosomes in yeast^47,48^, *Drosophila* (Vig2)^33^ and mammals (SERBP1)^31,33,49^. Based on these observations, we hypothesized that Habp4 and Dap1b/Dapl1/Dap play key roles in the prolonged ribosomal storage and/or translational repression in the egg. To test this hypothesis, we used CRISPR/Cas9-based mutagenesis to generate *habp4* knockout (*habp4*^*-/-*^) zebrafish lines (Extended Data Fig. 7a). *Habp4*^*-/-*^ mutants were viable and showed normal egg clutch sizes, fertility, embryo development and survival rates (Extended Data Fig. 7b-f). However, 1-3 hpf embryos derived from *habp4*^*-/-*^ parents contained about 30% less total RNA as well as 30% fewer monosomes when compared to 1-3 hpf wild-type siblings (Fig. 3e-g) despite having a similar average embryo size (Extended Data Fig. 7g). Given that 80-90% of the total cellular RNA is ribosomal RNA^50^, the reduction in total RNA content in *habp4*^*-/-*^ embryos is likely due to the loss of rRNA-containing ribosomes. In contrast, embryos from *habp4*^*-/-*^ and from wild-type parents displayed no significant difference in polysome levels at 1 hpf (Fig. 3g; Extended Data Fig. 7h), suggesting that Habp4 contributes to stabilizing 80S ribosomes but not to inhibiting translation in the zebrafish egg.

To analyze the physiological function of Dap1b/Dapl1 and Dap, we generated double mutant zebrafish lacking both Dap1b and Dap proteins (*dap*^*-/-*^, *dap1b*^*-/-*^) (Extended Data Fig. 7i). Adult *dap*^*-/-*^, *dap1b*^*-/-*^ mutant fish were obtained from a double heterozygous parent in-cross (*dap*^*+/-*^, *dap1b*^*+/-*^) at sub-Mendelian ratio compared to wild-type siblings (Fig. 3h), indicating a fitness defect. Though surviving *dap*^*-/-*^, *dap1b*^*-/-*^ adults appeared morphologically normal and showed normal egg clutch sizes, fertility, and embryo development and survival rates (Extended Data Fig. 7j-m), their progeny displayed a significantly higher polysome-to-monosome ratio at 1 hpf compared to 1 hpf embryos from wild-type fish (Fig. 3i, j; Extended Data Fig. 7n, o). Indeed, this ratio was similar to that observed for 2 hpf and 3 hpf embryos from wild-type parents (Fig. 3j), suggesting that Dap1b/Dap function as a translational inhibitor in dormant ribosomes of eggs and early embryos.

Dap1b/Dapl1 and Dap are small (∼15 kDa), unstructured proteins that are rich in basic amino acids and prolines (15%) (Fig. 4a; Extended Data Fig. 5). While invertebrates contain only one ortholog, vertebrates have evolved two paralogs (Fig. 4a, Extended Data Fig. 5). Despite displaying less than 50% overall sequence identity, all Dap1b/Dapl1 and Dap proteins share two highly conserved motifs at the N- (P-A-V-K-A-G-G-M/K-R) and C-terminus (I-Q/H-Q-P-R-K/R-x), which we found proximal to Rpl31 (Fig. 3c, d) and binding to the PET (Fig. 2a-c), respectively (Fig. 4a, Extended Data Fig. 5). To further investigate Dap1b/Dapl1’s and Dap’s function, we performed *in vitro* translation assays in rabbit reticulocyte lysate (RRL) in the presence of recombinant Dap1b/Dap proteins (Fig. 4b). Whereas recombinant zebrafish Dap or the negative control BSA did not affect the translational activity of RRL, recombinant zebrafish Dap1b repressed the translation of a luciferase mRNA (IC50: 4.7 µM) to a similar extent as the antimicrobial peptide Bac7 (IC50: 3.6 µM) (Fig. 4c), which binds to the PET and inhibits translation^16^. To investigate whether the observed difference in repressive activity between zebrafish Dap1b and Dap was generalizable to other species’ Dap1b/Dapl1 and Dap proteins, or whether it reflected a paralogue-specific mismatch between zebrafish Dap protein and mammalian ribosomes, we assessed ribosomal binding of *in vitro* translated flag-tagged *dap1b/dapl1* and *dap* mRNAs from various species (Fig. 4b). Consistent with a difference in repressive activity between Dap1b and Dap *in vitro*, Dap showed a weaker affinity for rabbit ribosomes than Dap1b/Dapl1 from three different vertebrate species (Fig. 4d, e; Extended Data Fig. 8a). Analysis of ancestral Dap1 proteins from invertebrates revealed that Dap1 from *C. elegans* and from the coral *Pocillopora* had an affinity for rabbit ribosomes in between that observed for vertebrate Dap and Dap1b proteins, while Dap from *D. melanogaster* did not bind to rabbit ribosomes (Fig. 4d, e; Extended Data Fig. 8a). Interestingly, we noticed that all rabbit ribosome-binding vertebrate Dap1b/Dapl1 proteins as well as the ancestral *Pocillopora* and *C. elegans* Dap1 proteins contain at least one additional residue at the C-terminus compared to non-binding vertebrate and *Drosophila* Dap proteins (Fig. 4a; Extended Data Fig. 5). However, extending zebrafish Dap or shortening zebrafish Dap1b by one amino acid did not have any significant effect on ribosome binding (Fig. 4f, g; Extended Data Fig. 8b). In contrast, mutating the conserved, hypusine-interacting Gln105 of zebrafish Dap1b (Fig. 2f) (as well as the corresponding Gln102 of zebrafish Dap) abolished binding of Dap1b (and Dap) to the ribosome (Fig. 4f, g; Extended Data Fig. 8b), thus establishing a key function of this invariant Glutamine in stabilizing Dap1b binding to the ribosome via its interaction with eIF5A. Together, these analyses revealed that Dap1b/Dapl1 but not Dap efficiently binds to mammalian ribosomes *in vitro* and is sufficient to inhibit translation.

**Fig. 4.**
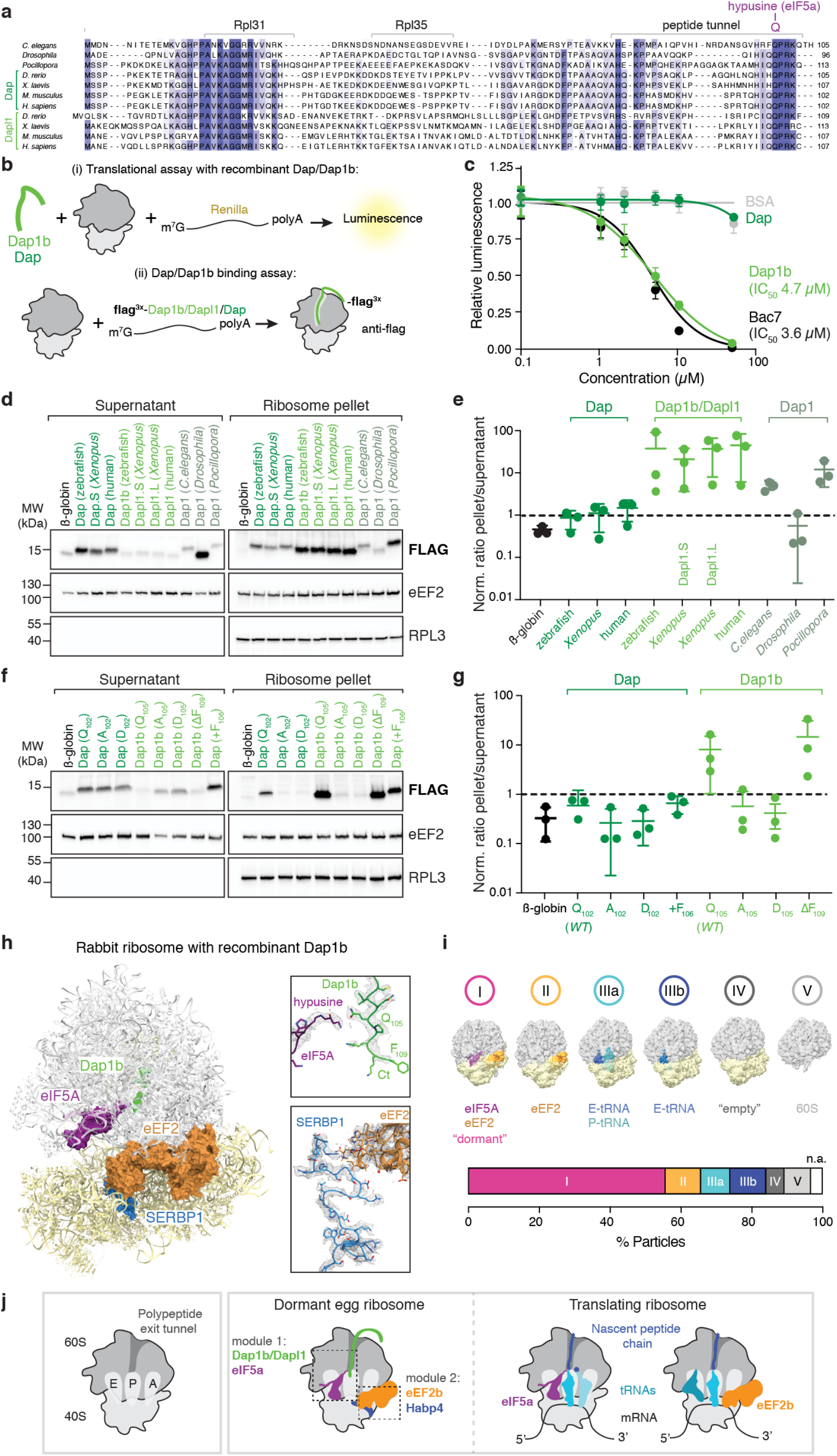
Dap1b/Dapl1 binding to mammalian ribosomes blocks translation and reconstitutes the egg-like ribosome-state *in vitro*. **a**, Alignment of Dap1b/Dapl1 (light green) and Dap (dark green) paralogs as well as ancestral Dap1 proteins from a subset of vertebrate and invertebrate species. Conserved regions are highlighted in shades of blue according to sequence identity (dark blue represents high concordance). A more extended alignment is shown in Extended Data Fig. 5a. **b**, Scheme of the *in vitro* translation assays performed in rabbit reticulocyte lysate (RRL). To investigate Dap1b/Dap activity (top), recombinant zebrafish Dap1b or Dap (or control proteins) and *in vitro* synthesized *Renilla luciferase* mRNA were added to RRL. Translation was assessed after 1.5 hours by measuring Luciferase activity. To assess Dap1b/Dapl1 and Dap binding to the ribosome (bottom), *in vitro* synthesized mRNAs encoding N-terminally tagged candidate proteins were incubated in RRL. Ribosomes were pelleted and Dap1b/Dapl1 and Dap levels were quantified by Western blot. **c**, IC50 analyses of the sufficiency of recombinant zebrafish Dap1b and Dap proteins to repress translation in RRL (see Fig. 4b, top). The antimicrobial peptide Bac7 and BSA were used as controls. For statistical analysis, four parameter logistic curve (best-fit solution, nonlinear regression-dynamic fitting) and normality tests (Kolmogorov–Smirnov) were used. **d**, Binding assays of *in vitro* translated proteins to rabbit ribosomes (see Fig. 4b, bottom). Western blots of ribosomal and supernatant fractions from binding assays with Flag-tagged β-globin (control) and Dap, Dap1b/Dapl1 and ancestral Dap1 from different species. eEF2 and RPL3 levels are used as readout of successful fractionation and for normalization. Blot of the total reaction is shown in Extended Data Fig. 8a. **e**, Quantification of Western blot signals detected in ribosome pellets compared to supernatants. Values are normalized to loading (eEF2/RPL3) and translation efficiency (FLAG signal of total lysate; Extended Data Fig. 8a) of the individual Flag-tagged factor mRNAs (n=3). **f**, Western blots of ribosomal and supernatant fractions from binding assays using Flag-tagged β-globin (control), as well as wild-type and mutant versions of zebrafish Dap and Dap1b. Mutant versions are indicated. Blot of the total reaction is shown in Extended Data Fig. 8b. **g**, Quantification of Western blot signals detected in ribosome pellets compared to supernatants. Values are normalized to loading (eEF2/RPL3) and translation efficiency (FLAG signal of total lysate; Extended Data Fig. 8b) of the individual Flag-tagged factor mRNAs (n=3). **h**, Structure of the ribosome from RRL supplemented with recombinant zebrafish Dap1b. Ribosome-associated factors are shown as surface representations. Densities (in mesh) for the two modules characteristic of dormant ribosomes (Dap1b-eIF5A and SERBP1-eEF2) are shown on the right. Critical amino acids are indicated. Ct, C-terminus. **i**, Distribution of ribosomal particles among the classes obtained after the analysis of particle heterogeneity in rabbit ribosomes supplemented with zebrafish Dap1b using cryoDRGN. Representative filtered density maps of the six major classes are shown on the top (see Extended Data Table 2; n.a refers to “non-assigned” particles). **j**, Scheme depicting the main features of the dormant ribosomes identified in this study. A cartoon of the important functional sites of the ribosome is shown on the left. Dormant ribosomes (middle cartoon) are associated with four factors which form two modules: the Habp4-eEF2 module occupies the mRNA channel and A site of the ribosome and interferes with mRNA and A-tRNA binding, and the Dap1b/Dapl1-eIF5A module blocks the polypeptide exit tunnel and the E/P sites. Two cartoons of a translating ribosome are shown on the right for comparison.

To assess whether Dap1b was inserted into the PET of mammalian ribosomes, and whether Dap1b addition to translational extracts indeed reconstitutes the dormant ribosome state, we performed cryo-EM on ribosomes isolated from an *in vitro* translation extract supplemented with 20 µM of recombinant zebrafish Dap1b and obtained a map at 2.3 Å average resolution (Fig. 4h; Extended Data Fig. 9, Extended Data Table 1). Notably, we were able to assign densities, not only for Dap1b in the PET (Fig. 4h, Extended Data Fig. 8c), but also for the other dormant ribosome factors, namely eIF5A, eEF2, and the Habp4 homolog SERBP1 (Fig. 4h, Extended Data Fig. 8c, d), which are present in RRL^31^. Rabbit ribosomes with bound Dap1b revealed additional interactions between eIF5A and RPL10A, which is part of the L1 stalk. We found that this interaction restrains the L1 stalk to a closed conformation, as previously reported for eIF5A-containing ribosomes^28^, which thus links the closed L1-stalk (with a locked E site) to the Dap1b-eIF5A module (Extended Data Fig. 8e, Extended Data Table 2). A 3D classification of these ribosomal particles using cryoDRGN^44^ revealed that, after addition of zebrafish Dap1b to RRL, 55.7 % of the particles contained both eIF5A and eEF2 (Fig. 4i (class I), Extended Data Fig. 8f, Extended Data Table 2), similarly to dormant ribosomes from 1 hpf zebrafish embryos and from *Xenopus* eggs (Fig. 2h, Extended Data Table 2). While several cryo-EM ribosome structures isolated from RRL have been published, none of them has reported the co-existence of eIF5A and eEF2^30,31^, which is consistent with Dap1b being causally linked to this dormant ribosome state. Together, our data reveal that zebrafish Dap1b can insert into the PET of mammalian ribosomes, which is sufficient to reconstitute the dormant ribosomal state *in vitro* and to efficiently block translation.

## Discussion

Here, we show that ribosomes are stored in zebrafish and *Xenopus* eggs in a newly identified dormant state, which can be reconstituted in mammalian ribosomes *in vitro*. This state is characterized by the binding of a conserved set of factors that occupy and thereby block functionally important sites of the ribosome (Fig. 4j). Our cryo-EM structures revealed two modules that co-exist on dormant ribosomes. One module is formed by Habp4-eEF2, in which binding of the egg-specific SERBP1-homolog Habp4 to the mRNA channel stabilizes eEF2 at the A site, which blocks mRNA and tRNAs from interacting with the ribosome. *In vivo* data reveal that Habp4 has a key role in stabilizing ribosomes in the egg (Fig. 3e-g), thus contributing to storing the large numbers of ribosomes that are needed for later embryogenesis. The other module is formed by Dap1b/Dapl1-eIF5a, in which insertion of Dap1b/Dapl1 into the PET mimics an arrested nascent peptide chain in the absence of bound mRNA or tRNAs and induces stable binding of eIF5a at the E/P site of the ribosome (Fig. 2a-c, Fig. 2e-f). Importantly, Dap1b binding to mammalian ribosomes is sufficient to induce the dormant ribosomal state *in vitro* (Fig. 4h, i) and to repress translational activity *in vivo* (Fig. 3i, j) and *in vitro* (Fig. 4c). Thus, the two modules act together to accomplish the specific needs of egg cells to (1) stably store ribosomes for prolonged periods of time, and (2) to repress translation. Importantly, repression of the translational machinery in the egg might also be important for the storage of maternal mRNAs since mRNA degradation has been shown to positively correlate with translation^51^. Moreover, having eIF5a and eEF2 stably bound to maternal ribosomes might thus preserve and protect these factors for their later use in embryogenesis, as eEF2 and eIF5a from *C. elegans* were shown to be prone to aggregate with aging^52^.

Our *in vivo* data suggest that Habp4, which occupies the mRNA entry channel, is important to preserve maternal ribosomes in the egg. Despite the lower rRNA and monosome content observed in early *habp4*^*-/-*^ embryos, they did not display any developmental defects under our standard growth conditions. This observation was surprising given that a reduction in ribosome levels had previously been shown to cause developmental defects (*minute* phenotypes in *Drosophila*^53^) and to alter cell differentiation (e.g. in embryonic stem cells^54^ and tumor formation in zebrafish^55^). We therefore speculate that ribosomes in zebrafish eggs are stored in excess since embryos can tolerate a reduction of about 30% of ribosomes. Maternal ribosomes could fulfil additional functions beyond protein synthesis, as they have been proposed to contribute to nucleotide homeostasis during animal development^56^. Other organisms with a smaller nutrient reservoir in eggs than zebrafish might be more dependent on the degradation of maternal ribosomes to obtain the nucleotides and amino acids necessary for some biosynthetic pathways. Therefore, Habp4 might be essential in organisms whose (i) maternal ribosomes must be stored for long periods of time and/or (ii) eggs do not contain a large nutrient reservoir.

Our study identified Dap1b/Dapl1 as a novel translational inhibitor bound to the PET of eukaryotic 80S ribosomes that is sufficient to reconstitute the egg ribosome state and can effectively block mammalian translation *in vitro* (Fig. 4c, Fig. 4h, i). This finding is remarkable since no endogenously made eukaryotic protein had been demonstrated before to be inserted into the PET of eukaryotic 80S ribosomes, leading to their repression. Several factors, including the antimicrobial peptides Bac7^16^ and Api137^57^, have been shown to be produced in eukaryotic cells as a defense against bacteria. They interact with the PET of bacterial ribosomes and repress bacterial translation (Extended Data Fig. 8g, h). In eukaryotes, only factors involved in ribosome biogenesis (e.g. Reh1, Rei1 and Nog1) have been observed to be inserted into the PET, yet not of mature ribosomes but during eukaryotic 60S ribosome biogenesis^58–61^ (Extended Data Fig. 8g, h). Given that *dap*^*-/-*^, *dap1b*^*-/-*^ mutants do not show any developmental phenotype during embryogenesis (Extended Data Fig. 7j, m) and that we do not observe an increase in the amount of polysomes in *dap*^*-/-*^, *dap1b*^*-/-*^ eggs (Extended Data Fig. 7o), we hypothesize that other inhibitory mechanisms, e.g. short polyadenine tails^13^ and inhibition of eIF4F^14,15^, can at least in part compensate for the loss of the dormant ribosomal state in the egg and keep translation low.

Two questions that will require future work is how Dap1b/Dapl1 is inserted into the PET of the ribosome, and how it is released after fertilization. Our *in vitro* data with recombinant Dap1b protein suggests that Dap1b is inserted into the PET post-translationally (Fig. 4c, h). Given the sequence conservation of the N- and C-terminus of Dap1b/Dapl1 and Dap proteins, and the presence of crosslinks between Dapl1’s and Dap’s N-terminus with Rpl31, we speculate that insertion may require both regions. Dissociation might be triggered by a single signaling event given that we observe a concerted release of all four factors during the first hours of embryogenesis (Fig. 1f). Since mammalian Dap1 has been shown to be phosphorylated in an mTOR-dependent manner^38^, this signaling event could be linked to the rise in mTOR signaling after fertilization^62^.

Zygotic ribosomes are first produced after the maternal-to-zygotic transition, an event that, in some species (e.g. in humans), can take place years after the mature oocyte is formed. While ribosome storage is remarkably different in the egg compared to any other so-far described ‘dormant’ ribosome state in somatic cells, it is intriguing to speculate that similar mechanisms may be used at other stages in the lifetime of animals that require prolonged periods of low metabolic activity, e.g. the diapause in *C. elegans* (dauer stage). The mechanism we describe here, namely having functionally important sites of the ribosome occupied by evolutionarily conserved factors, two of which are needed for resumption of translation later on, appears ideally suited to accomplish energy-preserving long-term storage of ribosomes in a variety of developmentally programmed and metabolically-induced contexts.

## Methods

### Zebrafish and *Xenopus* lines and husbandry

Zebrafish (*Danio rerio*) were raised according to standard protocols (28°C; 14/10 hour light/dark cycle). TLAB fish, generated by crossing zebrafish AB with the natural variant TL (Tupfel Longfin), served as wild-type zebrafish for all experiments. Zebrafish *dap, dap1b* and *habp4* mutants were generated as part of this study and are described below. Wild-type *Xenopus laevis* were obtained from NASCO (USA) and maintained in the IMP animal facility. All fish and *Xenopus* experiments were conducted according to Austrian and European guidelines for animal research and approved by local Austrian authorities (animal protocols for work with zebrafish: GZ 342445/2016/12 and MA 58-221180-2021-16; animal protocols for work with *Xenopus*: BMWFW-66.006/0012-WF/II/3b/2014, BMWFW-66.006/0003-WF/V/3b/2016).

### Generation of knockout fish lines

Zebrafish *dap, dap1b* and *habp4* mutants were generated by Cas9-mediated mutagenesis^63^. A pool of two guide-RNAs (sgRNAs) targeting the first exon of either *dap* (chr24:22071216-22103555), *dap1b* (chr9:52378482-52386733) or *habp4*/*zgc:103482* (chr21:20396891-20402554) were generated according to published protocols^63^ (see Extended Data Table 6 for oligos used for generating sgRNAs). Cas9 protein and sgRNA pools were co-injected into one-cell stage TLAB embryos. Putative founders were outcrossed to TLAB, and germline mutations were identified by a size difference in PCR amplicons (see Extended Data Table 6 for primers used for genotyping). Embryos from founders were raised to adulthood. Homozygous knockout fish were generated by crossing heterozygous fish. To generate *dap*^*-/-*^, *dap1b*^*-/-*^ double mutants, *dap*^*+/-*^ and *dap1b*^*+/-*^ fish were crossed, and double heterozygous mutants were genotyped and raised to adulthood. Double homozygous mutants (*dap*^*-/-*^, *dap1b*^*- /-*^) and wild-type (*dap*^*+/+*^, *dap1b*^*+/+*^) siblings were obtained from in-crossing double heterozygous mutants.

### Phenotypic characterization

Adult fish were crossed at least twice prior to phenotypic characterization. Females and males were kept at equal numbers per tank. Pictures of early development were taken at 0.5, 1, 2, 3, 6 and 24 hours postfertilization (hpf) on a ZEISS Stemi 508 stereo microscope with camera (2x magnification, FlyCapture2 software). To calculate the fertilization rate, the number of fertilized eggs was divided by the number of total eggs. Clutch size was calculated as the sum of fertilized and unfertilized eggs. Survival rate was calculated by dividing live larvae at 24 hpf by the number of fertilized eggs. The size of the embryos was measured as area (mm^2^) at 6 hpf in Fiji. Statistical analysis was performed in GraphPad Prism using Mann-Whitney test (for two groups, *p-value* < 0.05) or Kruskal-Wallis test (for multiple groups, *p-value* < 0.05).

### Polysome gradients

Wild-type and mutant embryos were dechorionated with pronase (1 mg/mL). 200 dechorionated embryos per genotype were lysed in 550 μL of lysis buffer (20 mM Tris-Cl pH 7.5, 30 mM MgCl_2_, 100 mM NaCl, 0.25 % Igepal-630 *(v/v*), 100 μg/mL cycloheximide, 0.5 mM dithiothreitol (DTT), and 1 mg/mL heparin). Embryo lysates were incubated for 10 min on ice and centrifuged for 10 min at 4°C with 20,000 xg. 200 μL of clarified lysates were loaded onto a continuous 10-50% *(w/v)* sucrose gradient prepared in TMS buffer (20 mM Tris-Cl pH 7.5, 5 mM MgCl_2_, 140 mM NaCl). Gradients were centrifuged in a SW40 Ti rotor (Beckman) at 4 °C and 35,000 rpm for 165 min. Polysome gradients were analyzed using a gradient station (BioComp) coupled to a Model Triax^™^ Flow Cell detector (FC-2). The precise location of the fractions along the UV-tracing was monitored using the Gradient profiler v1.25 (BioComp) software.

### RNA isolation

Total RNA was isolated from 10 homogenized zebrafish embryos per sample using the RNeasy Mini Kit (Qiagen). RNA concentration was measured on a spectrophotometer (DeNovix DS-11 FX+). Mutant RNA concentration was normalized to wild-type RNA concentration for each experiment.

### Analysis of translation efficiency

To calculate translational efficiencies over the time-course of embryogenesis, published PolyA+ RNASeq data^24^ (GSE32898) and ribosome profiling data^23^ (GSE46512) was pre-processed according to standard bioinformatic procedures. For ribosome profiling data (ribo-seq), 3’ adaptor sequences were removed using bbduk from bbmap v38.26, and trimmed reads were mapped to a set of rRNAs and abundant sequences downloaded from the Illumina iGenome using bowtie2. The remaining reads were aligned to the zebrafish reference genome GRCz10 using tophat2. For RNA-seq analysis, adaptor sequences were removed with bbduk from bbmap v38.26, and abundant reads that mapped to abundant sequences using bowtie2 were removed, similar to ribo-seq reads. Remaining reads were mapped with tophat2 to the zebrafish transcriptome using the GRCz10 genome assembly and the Ensembl 82 transcriptome release.

For ribo-seq and RNA-seq, FPKM values for the coding sequence (CDS) were calculated for transcripts with an annotated start and stop codon. FPKM values were defined as read counts over CDS / (CDS length * total read count) * 10^9^. Translational efficiencies were obtained by dividing FPKM of ribo-seq by FPKM of RNA-seq. TE values + 0.1 were used for plotting.

### Sample collection for ribosome isolation

The evening prior to the zebrafish egg collections, male and female zebrafish were separated in breeding cages. To collect mature, un-activated eggs, female zebrafish were anesthetized in the morning using 0.1 *% (w/v)* Tricaine (25x stock solution in dH2O, buffered to pH 7-7.5 with 1 M Tris pH 9.0). After being gently dried on a paper towel, the female was transferred to a dry petri dish, and eggs were carefully expelled from the female by applying mild pressure on the fish belly with a finger and stroking from anterior to posterior. The eggs were separated from the female using a small paintbrush, and the female was transferred back to the breeding cage filled with fish water for recovery. To prevent activation, eggs were kept in sorting medium (Leibovitz’s medium, 0.5 % BSA, pH 9)^64^ at room temperature (RT). In the case of zebrafish embryos (1,000 embryos per sample for mass spectrometry (MS) and electron cryo-microscopy (Cryo-EM), and 5,000 embryos per sample for crosslinking-MS), eggs were collected in Petri dishes with blue water, which consist of fish water, 0.025 % *(v/v)* Instant Ocean salts (Aquarium Systems, 218035), and 0.0001 % *(v/v)* methylene blue (Sigma-Aldrich, M9140), pH 7. Embryos were incubated at 28°C, collected at the desired time points (1 hpf and 6 hpf), and incubated with 1 mg/mL of pronase for 5 min at RT for dechorionation.

To obtain samples from *Xenopus*, egg collection and *in vitro* fertilization were performed following a previously described protocol^65^. In brief, sexually matured wild-type females were primed with 50 IU of pregnant mare serum gonadotropin (PMSG, ProSpec, HOR-272) one week before the experiments. At the evening before egg collection, PMSG-primed females were injected with 500 IU of human chorionic gonadotropin (hCG, Sigma-Aldrich, CG10). Freshly laid eggs were collected and chemically dejellied in 2% *(w/v)* L-cysteine/1xMMR (Marc’s Modified Ringer) pH 7.8 before ribosome extraction. For the 24 hpf sample, freshly laid eggs were *in vitro* fertilized with wild-type sperm and dejellied in 2% *(w/v)* L-cysteine/0.1xMMR pH 7.8 at the 1-cell stage. Embryos were allowed to develop at RT for 24 h before ribosome extraction.

### Ribosome isolation

Samples were washed in 2 volumes of lysis buffer, containing 20 mM HEPES-KOH pH 7.4, 150 mM KCl, 10 mM MgCl_2_, 0.5 mM DTT, rRNasin (Promega), 0.25% RNaseOUT *(v/v)* (ThermoFisher), 0.25% SUPERaseIN *(v/v)* (ThermoFisher), 0.25% Igepal *(v/v)*, and cOmpleteTM-EDTA-free protease inhibitor (Roche). Embryos were lysed in 2 mL (for 1,000 embryos) or 5 mL (for 5,000 embryos) of lysis buffer using a pre-cooled Dounce homogenizer. 100 μg/mL of cycloheximide was added during lysis. Following incubation on ice for 5 min, the supernatant was cleared by centrifugation at 20,000 xg for 10 min at 4°C. Ribosome isolation was adapted from ^66^. Briefly, the clarified supernatant was loaded onto a 30% *(w/v*) sucrose cushion prepared in buffer A (20 mM Tris-Cl pH 7.5, 2 mM Mg(OAc)_2_ and 150 mM KCl) and centrifuged at 116,000 xg for 5 h at 4°C in a TL100.3 rotor (Beckman). The ribosome pellet was resuspended by orbital shaking (1 h at 4°C) in buffer B, consisting of 20 mM Tris-Cl pH 7.5, 6 mM Mg(OAc)_2_, 150 mM KCl, 6.8% sucrose *(w/v*), 1 mM DTT, 0.25% RNasin Plus *(v/v)* (ThermoFisher), and 0.25% RNase Inhibitor *(v/v)*, (Promega). Resuspended ribosomes were loaded onto a 15-30% *(w/v)* sucrose gradient prepared in buffer C (100 mM KCl, 5 mM Mg(OAc)_2_, 20 mM HEPES-KOH pH 7.6, 1 mM DTT, and 10 mM NH_4_Cl) and centrifuged for 16 h at 4°C with 18,000 rpm in an SW60Ti rotor (Beckman). Gradients were hand-fractionated and ribosome concentrations were determined by absorbance at 260 nm (A_260_) and SDS-PAGE.

To isolate ribosomes from rabbit reticulocyte lysates (RRL) (Green Hectares; composition and preparation of the lysate are described in the section “*In vitro* translation assays” below), RRL was supplemented with 20 µM recombinant zebrafish Dap1b protein and incubated for 1 h at 30°C. A one-step centrifugation of 1 mL of *in vitro* translation reaction was performed over 3 mL of a 30% *(w/v)* sucrose cushion prepared in RNC buffer (50 mM HEPES-KOH pH 7.4, 100 mM KOAc, 5 mM Mg(OAc)_2_, and 1 mM DTT) in a TLA100.3 rotor (Beckman) at 116,000 xg for 5 h. The ribosome pellet was resuspended in 120 mL of RNC buffer.

For MS and cryo-EM, sucrose was omitted from the gradient fractions by loading the resuspended ribosome solution on a Zeba™ Spin Desalting Columns (7K MWCO).

### Digest of total cell lysates and ribosomes for mass spectrometry (MS)

Total cell lysates and isolated ribosomes from zebrafish eggs and embryos and from *Xenopus* eggs were either directly denatured in 8 M urea in 100 mM ammoniumbicarbonate (ABC) buffer or acetone precipitated before being dissolved in 8 M urea in 100 mM ABC. DTT was added to a final concentration of 10 mM and the sample was incubated 1 h at 37°C. Alkylation was performed by adding iodoacetamide (IAA) to a final concentration of 20 mM and incubating for 30 min at RT in the dark. The reaction was quenched by addition of 5 mM DTT and incubated again 30 min at RT.

Samples were diluted to 6 M urea with 100 mM ABC followed by addition of Lys-C (FUJIFILM Wako Pure Chemical Corporation) at a ratio of 1:100 and incubation at 37°C for 2 h. The samples were diluted to 2 M urea with 100 mM ABC. Trypsin (Promega, Trypsin Gold) was added at a ratio of 1:100 and incubated at 37°C overnight. 500 ng of each sample were analyzed by liquid chromatography-mass spectrometry (LC-MS/MS).

### Crosslinking of purified ribosomes and digest for mass spectrometry (MS)

For elution of zebrafish 1 hpf and *Xenopus* egg ribosome fractions used in crosslinking-MS experiments, buffer C was adjusted to 20 mM HEPES-KOH pH 7.6, 100 mM KCl, and 5 mM Mg(OAc)_2_. 0.5 mM of disuccinimidyl sulfoxide (DSSO) from a 5 mM stock solution (in dimethyl sulfoxide, DMSO) was added to 10 µL of 500 nM ribosomes and incubated for 45 min at RT. The reaction was quenched by adding Tris-Cl pH 7.5 to a final concentration of 100 mM and incubated for 15 min at RT. The band pattern of the crosslinking reaction was analyzed by SDS-PAGE.

DSSO crosslinked ribosomes were denatured in 8 M urea, 100 mM ABC followed by reduction, alkylation and proteolytic digest using Lys-C and trypsin (both added 1:20) as described above. Digests were acidified to 1% TFA and desalted using Oasis HLB Sorbent (Oasis HLB Microelution plate, Waters) according to the manufacturer’s description. Peptides were eluted with 2 × 30 μL 70% acetonitrile (ACN), 0.1% TFA and the sample was concentrated under reduced pressure to 20 μL. The samples were supplemented with 5% DMSO. To enrich for crosslinked peptides, the samples were fractionated by size-exclusion chromatography (SEC) on a TSKgel SuperSW2000 column (300 mm × 4.5 mm × 4 μm, Tosoh Bioscience), which was operated at 200 μL/min in 30% ACN, 0.1% TFA. Fractions were collected every minute, and ACN was removed under reduced pressure. DMSO was again added to 5%.

### LC-MS/MS

Generated peptides were analyzed on a nano-reversed phase HPLC (RSLC nano system, ThermoFisher) coupled to a Q Exactive HF-X mass spectrometer (ThermoFisher), equipped with a Nanospray Flex ion source (ThermoFisher).

Peptides were loaded onto a trap column (PepMap C18, 5 mm × 300 μm ID, 5 μm particles, 100 Å pore size) at a flow rate of 25 μL/min using 0.1% TFA as mobile phase. After 10 min, the trap column was switched in line with the analytical column (PepMap C18, 500 mm × 75 μm ID, 2 μm, 100 Å, ThermoFisher), which was operated at a flowrate of 230 nl/min at 30°C. For separation a solvent gradient was applied, starting with 98% buffer A (0.1% formic acid in water) and 2% buffer B (80% ACN, 0.1% formic acid), followed by an increase to 35% buffer B over the next 180 or 360 min for digested lysates and ribosomes. For unfractionated and SEC-enriched crosslinked samples, a gradient to 40% buffer B in 180 or 120 min, respectively, was used. This was followed by a steep gradient to 90% buffer B in 5 min, staying there for five min and decreasing to 2% buffer B in another 5 min.

For the digested lysates and ribosomes, the mass spectrometer was operated in data-dependent mode, using a full scan (m/z range 375-1500, resolution of 60,000, target value 1E6) followed by MS/MS scans of the 10 most abundant ions. MS/MS spectra were acquired using a normalized collision energy of 28, isolation width of 1.4 m/z, resolution of 30,000, maximum fill time of 105 ms and a target value of 1E5. Precursor ions selected for fragmentation (excluding charge state 1, 7, 8, >8) were put on a dynamic exclusion list for 60 s. Additionally, the minimum AGC target was set to 5E3 and intensity threshold was calculated to be 4.8E4. The peptide match feature was set to preferred, and the exclude isotopes feature was enabled.

For crosslinked samples, the mass spectrometer was operated in data-dependent mode, using a full scan (m/z range 350-1600, resolution of 120,000 and a target value of 1E6) followed by MS/MS scans of the 15 most abundant ions. MS/MS spectra were acquired using a stepped normalized collision energy of 27+/-6, isolation width of 1 m/z, resolution of 30,000, maximum fill time of 150 ms and a target value of 5E4. Precursor ions selected for fragmentation (excluding charge state 1, 2, >7) were put on a dynamic exclusion list for 30 s. Additionally, the minimum AGC target was set to 5E3, and intensity threshold was calculated to be 3.3E4. The peptide match feature was set to preferred, and the exclude isotopes feature was enabled.

### Analysis of MS data

For peptide identification, the RAW-files were loaded into Proteome Discoverer (v2.5.0.402, ThermoFisher). All hereby created MS/MS spectra were searched using MSAmanda v2.5.0.16129, Engine v2.0.0.16129^67^. For the first step search, the RAW-files were searched against a combined zebrafish sequence database (Danio_rerio GRCz11) and Uniprot downloaded on 21^st^ March 2019 (58,524 sequences; 34,079,443 residues), and against the *Xenopus laevis* Uniprot proteome UP000186698. All RAW-files were supplemented with common contaminants using the following search parameters: (i) the peptide mass tolerance was set to ±5 ppm and the fragment mass tolerance to ±8 ppm, and (ii) the maximal number of missed cleavages was set to 2 using tryptic enzymatic specificity. The result was filtered to 1% FDR on protein level using Percolator algorithm^68^ integrated in Proteome Discoverer. A sub-database was generated for further processing. For the second step, the RAW-files were searched against the created sub-database. Oxidation on methionine, deamidation on asparagine and glutamine, phosphorylation on serine, threonine and tyrosine; iodoacetamide derivative on cysteine, beta-methylthiolation on cysteine and carbamylation on lysine were set as variable modifications. Monoisotopic masses were searched within unrestricted protein masses for tryptic enzymatic specificity. The peptide mass tolerance was set to ±5 ppm and the fragment mass tolerance to ±8 ppm. The maximal number of missed cleavages was set to 2. The result was filtered to 1% FDR on peptide level using Percolator algorithm integrated in Proteome Discoverer. The localization of the post-translational modification sites within the peptides was performed with the tool ptmRS, based on the tool phosphoRS^69^. Peptide areas have been quantified using in-house-developed tool apQuant^70^. Proteins were quantified by summing unique and razor peptides and applying the intensity-based absolute quantification (iBAQ) calculation. Normalization of protein abundances was performed on the sum of areas of all identified ribosomal proteins. Statistical significance of differentially expressed proteins was determined using limma^71^.

### Analysis of crosslinking-MS data

Data analysis was performed within Proteome Discoverer v2.5.0.402 using MS Annika (v1.0.18345)^72^. The workflow tree consisted of the MS Annika Detector node (MS tolerance 10 ppm, crosslink modification: DSSO +185.004 Da at lysine and at protein N-termini, diagnostic ions: 138.0911; 155.179; 170.0634; 187.0900, crosslink modification addition: 18.010565, doublet pair selection in combined mode) followed by MS Annika Search (full tryptic digest, 5/10 ppm peptide/fragment mass tolerance, maximal 3 missed cleavages, carbamidomethyl +57.021 Da at cysteine as static and oxidation +15.995 Da at methionine as dynamic modification) and completed with MS Annika Validator (1% FDR cutoff at CSM and crosslink level, separate Intra/Inter-link FDR set to false). The search was performed against a database generated from a MS analysis of a digested non-crosslinked zebrafish (containing 1,092 proteins) or *Xenopus* (containing 1,173 proteins) ribosome sample, respectively. To generate these specific databases the two non-crosslinked samples were searched using MS Amanda against the zebrafish Uniprot reference proteome UP000000437 (downloaded 2020-09-15, 46,847 sequences, 24,556,292 residues) or *Xenopus laevis* Uniprot proteome UP000186698 (downloaded 2020-01-07, 43,235 sequences, 19,251,456 residues), both supplemented with common contaminants. Search parameters were set as described for time course experiments but using 5/10 ppm peptide/fragment mass tolerance and maximum 3 missed cleavages. Carbamidomethylation on cysteine was set as a fixed modification, oxidation on methionine, deamidation on asparagine and glutamine and acetylation on the protein N-terminus were set as variable modifications.

To validate the quality of the chosen search strategy, an additional crosslink search was performed using equal settings but a database file containing sequences aligned to the dormant ribosome structures of *Xenopus* and zebrafish. The resulting crosslinks at 1% FDR were plotted onto the structure using ChimeraX v1.1^73^, and the distance between connected amino acids (Cα-Cα distance) was measured. Crosslinks including Cα residues missing in the structure file were removed, yielding 360 and 178 crosslinks aligned to the zebrafish and *Xenopus* structures, respectively.

### DNA constructs and mRNA synthesis

*dap, dap1b, habp4, renilla luciferase* and zebrafish *β-globin* constructs for *in vitro* translation in rabbit reticulocyte lysate (RRL) (Green Hectares) were cloned into pCDNA3.1 by Gibson assembly^74^. Zebrafish coding sequences were amplified from cDNA. *Dap, dapl1* and *dap1* coding sequences from other species were ordered as Gblocks from IDT (amino acid sequences are shown in Extended Data Fig. 5). Capped mRNAs of all cloned constructs were synthesized with the Sp6 mMessage Machine kit (Ambion) according to the manufacturer’s protocol, using linearized plasmids as templates.

### Recombinant protein expression and purifications

Zebrafish *dap1b* and *dap* coding sequences were codon optimized for *E. coli*, cloned into a vector providing an N-terminal 6xHis tag followed by a 3C protease-cleavage site, and transformed into BL21(DE3) *E. coli* cells. For Dap1b expression, cells were grown in LB at 37°C until reaching an OD_600_ of 0.7, induced with 0.5 mM IPTG, and grown overnight at 18°C. For Dap expression, bacterial cultures were grown on terrific broth medium supplemented with 1.5% *(w/v)* lactose for 1 h at 37°C, and for 23 h at 18°C. Cells were pelleted, resuspended in lysis buffer (50 mM Tris-Cl pH 8.0, 300 mM NaCl, cOmplete Protease Inhibitor Cocktail [Merck]) and lysed by sonication (19 mm probe, 4 min: pulse on 1 s, pulse off 2 s, 60% amplitude). The intracellular soluble fraction was isolated by centrifugation at 20,000 xg, 30 min, 4°C. Imidazole was added to a final concentration of 20 mM. The lysate was loaded onto a 5 mL HisTrap FF column (Merck), washed with buffer A (50 mM Tris-Cl pH 8.0, 300 mM NaCl), and eluted with buffer A supplemented with 500 mM imidazole. 1 mg of 3C protease was added to the elution fraction during dialysis overnight against buffer A. The cleaved protein was loaded onto a 5 mL HisTrap FF column (Merck); the flow-through, containing the cleaved protein, was concentrated (Vivaspin 20, MWCO 5 kDa) and loaded onto a Superdex 75 26/60 column (GE Life Sciences) equilibrated with SEC buffer (20 mM HEPES-KOH pH 7.5, 150 mM KCl).

### *In vitro* translation (IVT) assays

Rabbit reticulocyte lysate (RRL; Green Hectares) was optimized for translation as previously described^75,76^. The final reaction contained 35% *(v/v*) RRL, 20 mM HEPES-KOH pH 7.4, 10 mM KOH, 50 mM KOAc, 1 mM ATP, 1 mM GTP, 12 mM creatine phosphate, 0.1 mg/mL tRNAs (Merck), 40 μg/mL creatine kinase, 2 mM MgCl_2_, 1 mM reduced glutathione, 0.3 mM spermidine, and 40 μM of each of the 20 amino acids. 1 μg of mRNA was added to a total volume of 30 μL of RRL. To compare translation of different factors, equimolar amounts of mRNA were added per reaction. Reactions were performed at 30°C for 60 min (unless stated otherwise). To assess ribosome binding, 250 μL of the reaction was loaded onto a 30% *(w/v)* sucrose cushion in ribosome-stabilizing RNC buffer (50 mM HEPES-KOH pH 7.4, 100 mM KOAc, 5 mM Mg(OAc)_2_, and 1 mM DTT), and centrifuged for 5 h with 116,000 xg at 4°C in a TLA100.3 rotor (Beckman). Binding to the different fractions was assessed by Western blot. To test for translation inhibition of recombinant Dap and Dap1b proteins, 0.1, 1, 2, 5, 10 or 50 μM recombinant protein in protein buffer (20 mM HEPES-KOH pH 7.4, 150 mM KCl) and 1 μg *Renilla* mRNA were added to RRL to a total volume of 25 μL. Bac7(1-35) synthetic peptide (1-RRIRPRPPRLPRPRPRPLPFPRPGPRPIPRPLPFP-35) was used as a positive control. BSA served as negative control. Reactions were performed at 30°C for 90 min. *Renilla* luciferase activity (Promega, E1960) was measured in duplicates on a Synergy H1 plate reader (BioTek). Samples were normalized to buffer control containing 1 μg of *renilla luciferase* mRNA. Relative IC50 was calculated in GraphPad Prism using nonlinear regression and the equation [Inhibitor] vs. response - Variable slope (four parameters).

### Western blotting

Western blotting was performed following standard protocols. In brief, protein samples were boiled for 5 min in 1x Laemmli sample buffer and separated on an SDS-PAGE using Mini-PROTEAN® TGXTM Precast Protein Gels or 10–20% Mini-PROTEAN® Tris-Tricine Gels (Bio-Rad). Blotting was performed on a nitrocellulose membrane (GE Healthcare) using a wet-blot system (BioRad). The following antibodies were used: anti-FLAG (mouse, 1:1000, Sigma-Aldrich F1804), anti-eEF2 (rabbit, 1:1000, Proteintech, 20107-1-AP) and anti-RPL3 (rabbit, 1:1000, GeneTex, GTX124464). The chemiluminescent signal was quantified using ImageJ 1.8.0_172. To quantify ribosome binding of FLAG-tagged proteins after *in vitro* translation, signal intensities were simultaneously measured for FLAG, RPL3 and eEF2 for all three membranes (total reaction, supernatant, ribosome pellet). The ratio between the FLAG signal in the supernatant and the ribosome pellet was normalized to the total amount translated (FLAG signal in the total reaction) and to sample loading (RPL3 and eEF2 signals).

### Grid preparation for cryo-EM

A 2 nm thick continuous carbon film was produced on an Auto306 high vacuum evaporator (Boc Edwards). This film was floated onto Quantifoil Cu 3.5/1, 200 mesh grids, and grids were dried. Grids were glow-discharged for 1 min in a SCD005 sputter coater (Bal-Tec) at ∼20 mA. 4 µl of sample (at a concentration of 200 ng/µl of RNA, which corresponds to ∼ 100 nM ribosomes) were applied to the grid and incubated for 60 s at 70% humidity and 4°C in a Leica EM GP. Subsequently, grids were blotted for 2 s using the proximity sensor and plunge-frozen in liquid ethane at –180°C.

### Cryo-EM data collection

All grids were screened on a Glacios TEM (ThermoFisher) to check for particle distribution and grid quality. Grids that past the evaluation were recorded on a Titan Krios microscope equipped either with a Falcon 3EC or a K3 detector. A pixel size of about 1 Å/px was chosen, and micrographs with a total dose of about 40 e/Å2 fractionated in 39 frames were collected with a target defocus between –1 and –2.5 µm.

### Electron microscopy data processing

Most data processing was performed in Cryosparc version 2.16.0^77^. Micrographs were motion corrected and dose weighted using Patch motion correction. CTF (Contrast Transfer Function) parameters were determined using Patch CTF. About 500 particles were manually selected and 2D classified to create templates. Automated particle picking was performed using these templates. Picking was manually inspected, and a threshold that excludes low signal-to-noise and high signal-to-dirt false positive picks and minimizes the false negative picks was chosen. The remaining particles were extracted with a box size between 440 and 512 and subjected to 2D classification. Classes that showed clear ribosomal densities were selected and subjected to heterogeneous ab-initio model generation. Subsequently, the remaining particles were 3D refined to generate a consensus model. Afterwards, optics groups were assigned to the individual particles depending on the applied beam shift during recording. Global and local CTF refinements were then performed, and all particles were refined using non-uniform refinement. To generate masks for local refinements, the density was filtered to 20 Å, and sub-densities of the entire 40S subunit and the head region were generated using the Volume eraser tool in UCSF Chimera 1.13.1^78^. These densities were binarized, extended by 7 pixels and another 7 pixels as soft edge. Locally refined maps using the corresponding masks were obtained in Cryosparc, and composite maps were generated with the *vop maximum* command in UCSF Chimera 1.13.1^78^.

To analyze particle dynamics and heterogeneity, the consensus refined particles were subjected to CryoDRGN analysis^44^. Due to the large pixel box size of our data sets, the dimensions of the ribosome particles were reduced to allow for feasible computation. The particles were down-sampled to a box size of 128 or 256, and two iterative rounds of the CryoDRGN VAE were trained using a network with 3 decoder and 3 encoder layers with 1024 as outer dimensions and a bottleneck latent dimension of 8. The resulting z values were classified using *kmeans* clustering and gaussian mixture models and plotted using a UMAP approach.

### Model building and refinement

The molecular models of the 1 hpf and 6 hpf ribosomes from zebrafish were built using PDB-4UG0^79^ as an initial model for core ribosomal proteins and rRNA. In the case of the 1 hpf zebrafish ribosome, additional factors were modeled from previously published structures: eIF5A from PDB-5DAT, and eEF2 and Habp4 (SERBP1 paralog) from PDB-6MTE^31^. All chains were first rigid body-fitted in Coot 0.8.9^80,81^ and manually mutated to the zebrafish sequences, taking MS data from purified zebrafish ribosomes (protein identity and abundance) into consideration. In the case of Habp4, another protein (Serbp1a, Uniprot ID F1Q5Q3) could also fit into the 1 hpf ribosome density. Given that Habp4 was more abundant than Serbp1a in 1 hpf zebrafish ribosomes according to MS, Habp4 was modeled.

The dormant ribosome structure from *Xenopus* eggs was built based on the 60S subunit of the 1 hpf zebrafish ribosome, and the 40S subunit of the 6 hpf zebrafish ribosome, given that both 40S subunits showed the same swiveled conformation. Protein and rRNA molecules from zebrafish were mutated to the corresponding sequences in *Xenopus*, taking into consideration the MS data from purified *Xenopus* ribosomes. Dapl1.S was built *de novo* in Coot and then used as a template to build Dap1b in the zebrafish 1 hpf ribosome. Although two Dap proteins (Dapl1.S and Dapl1.L in *Xenopus*, and Dap1b and Dap in zebrafish) could fit into the corresponding densities, Dapl1.S and Dap1b were modeled according to gene expression data (Extended Data Fig. 6) and protein abundance (measured by MS).

The rabbit ribosome model with recombinant zebrafish Dap1b was built using 6MTE^31^ as an initial model for core ribosomal proteins, rRNA, eEF2 and SERBP1. eIF5A was modeled using PDB-5DAT as reference. All models were manually adjusted to fit the observed density in Coot 0.8.9^80,81^ and real-space-refined using Phenix 1.17.1^82^. The 40S subunit of the rabbit ribosome was only submitted to rigid body fitting in Phenix. Working files were saved in PDB and the final models were converted to mmCIF using PDB_extract^83^. Cryo-EM densities and models were visualized using UCSF ChimeraX version 0.91^73^.

## Data availability

Cryo-EM maps and molecular models have been deposited in the Protein Data Bank (PDB) with IDs 7OYA (zebrafish 1 hpf), 7OYB (zebrafish 6 hpf), 7OYC (*Xenopus* eggs) and 7OYD (rabbit ribosome with zebrafish Dap1b), and in the Electron Microscopy Data Bank with accession codes EMD-13111 (zebrafish 1 hpf), EMD-13112 (zebrafish 6 hpf), EMD-13113 (*Xenopus* egg ribosome) and EMD-13114 (rabbit ribosome with zebrafish Dap1b). Raw micrographs, particle stacks and CryoDRGN models were deposited at the EMPIAR Database. Mass spectrometry proteomics data have been deposited to the ProteomeXchange Consortium via the PRIDE^84^ partner repository with the dataset identifier PXD026866.

## Acknowledgements

We thank L.E. Cabrera-Quio for ribosome profiling data analysis; K.R. Gert for providing non-activated zebrafish eggs; the Mass Spectrometry Facility at the Vienna BioCenter Core Facilities GmbH (VBCF), in particular S. Opravil for the processing of our samples, and R. Imre and G. Dürnberger for their help with data analysis; the VBCF Protein Technologies facility, in particular A. Lehner, for the expression and purification of recombinant Dap1b and Dap proteins; M. Madalinski for synthesizing the Bac7 peptide; the VBCF Electron Microscopy Facility for the support and maintenance of the facilities; H. Stark (MPI BPC) for providing additional Krios measurement time; the IMP animal facility personnel, in particular J. König and F. Ecker, for their excellent care of fish and *Xenopus*; M. Binner, J. Kiraly and A. Bandura for their help with genotyping; E. Tanaka and T. Clausen for funding T.-Y.L. and A.M., respectively; the entire Pauli group for valuable discussions on the project; the VBC RNA Salon and the RNA Deco-SFB community for providing useful feedback and suggestions; A. Andersen (Life Science Editors), C. Plaschka, A. Stark, J. Brennecke, A. Carter, F. Peske, U. Hohmann, E. Calo, A. Shah and A. Blaha for critical reading and valuable feedback on the manuscript. This work was supported by the Institute of Molecular Pathology (IMP), which receives institutional funding from Boehringer Ingelheim and the Austrian Research Promotion Agency (Headquarter grant FFG-852936), and funding from the FWF START program (Y 1031-B28 to A.P.), the Human Frontier Science Program (HFSP) Career Development Award (CDA00066/2015 to A.P.), the SFB RNA-Deco (project number F 80 to A.P.) and EMBO-YIP funds (to A.P.). L.L.-O. was supported by an SNF Early Postdoc Mobility fellowship (P2GEP3_191204) and an EMBO long-term fellowship (ALTF 1165-2019); K.R.G. was supported by a DOC Fellowship from the Austrian Academy of Sciences. We acknowledge the cryo-electron microscopy facility CEITEC MU of CIISB, Instruct-CZ Centre, supported by MEYS CR (LM2018127), and Diamond Light Source for granting us access and support at the cryo-EM facilities at the UK’s national Electron Bio-imaging Centre (eBIC) under proposal EM BI25222, funded by the Wellcome Trust, MRC and BBRSC. For the purpose of Open Access, the authors have applied a CC BY public copyright license to any Author Accepted Manuscript (AAM) version arising from this submission.

## Author contributions

F.L., L.L.-O., D.H. and A.P. conceived the study; F.L. performed most experiments with the help of C.P.; L.L.-O. obtained the final cryo-EM maps and modeled all ribosomes with contributions of F.L., A.M. and D.H. and help from S.K.; I.G. prepared and screened grids, collected EM data, and together with F.L. and D.H., contributed to the initial cryo-EM data processing with help from K.B.; M.M. and E.R. performed and analyzed the crosslinking sample preparation, and K.M. supervised mass-spectrometry experiments and analyzed the crosslinking samples; T.-Y.L. provided *Xenopus* eggs and 24 hpf embryos; D.H. performed the cryoDRGN analysis and F.L. and L.L-O. compared the resulting maps; A.P. and D.H. supervised the project; L.L.-O., F.L., and A.P. wrote the manuscript with input from all authors.

## Competing interests

The authors declare no competing interests.

## Materials & Correspondence

Correspondence and requests for materials should be addressed to A.P.

## Extended Data Figures & Tables

**Extended Data Fig. 1.**
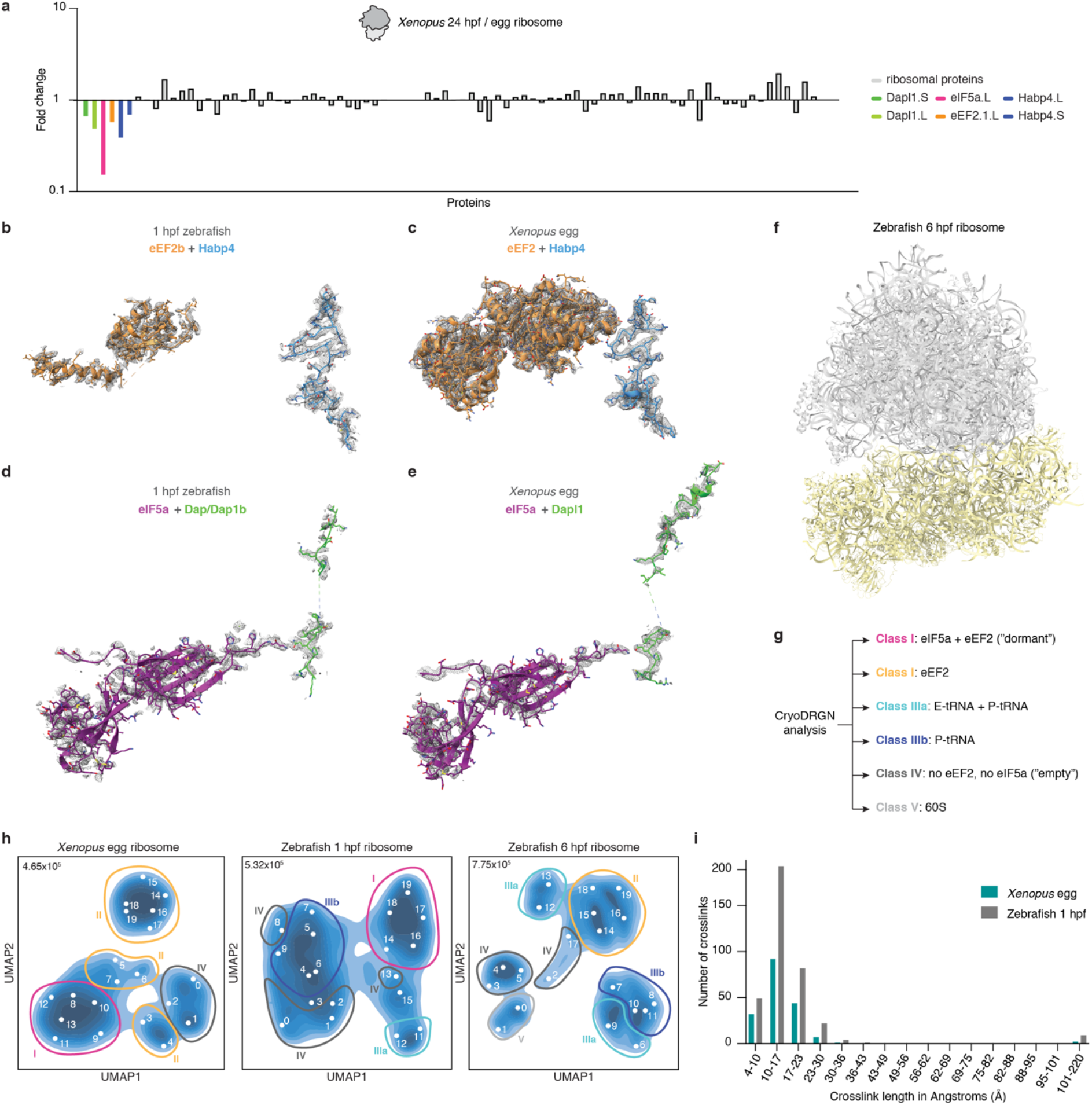
Characterization of the dormant ribosome state in zebrafish 1 hpf and *Xenopus* egg ribosomes. **a**, Fold change of ribosome-associated factors and core ribosomal proteins after comparing mass spectrometry data of purified ribosomes from unfertilized *Xenopus* eggs and 24 hpf larvae (stage 14) (n=1). **b-e**, Densities of the two modules, namely Habp4-eEF2b/eEF2 (b-c), and Dap1b/Dapl1-eIF5a (d-e), that are characteristic for dormant ribosomes in zebrafish 1 hpf embryos and *Xenopus* eggs. **f**, Overview of the ribosome structure isolated from 6 hpf zebrafish embryos lacking ribosome-associated factors. **g**, Classes assigned to the volumes obtained after the analysis of ribosomal particle heterogeneity using cryoDRGN. **h**, Latent space representations of ribosomal particles from *Xenopus* eggs (left), 1 hpf zebrafish embryos (middle) and 6 hpf zebrafish embryos (right) as UMAP embeddings after training a cryoDRGN latent variable model. Classes are depicted with circles in Roman numbers, map volumes are indicated with Arabic numbers. Total particle numbers are shown on the top left of each graph. **i**, Cα-Cα distance distribution of the DSSO-induced crosslinks identified in *Xenopus* egg and zebrafish 1 hpf ribosomes.

**Extended Data Fig. 2.**
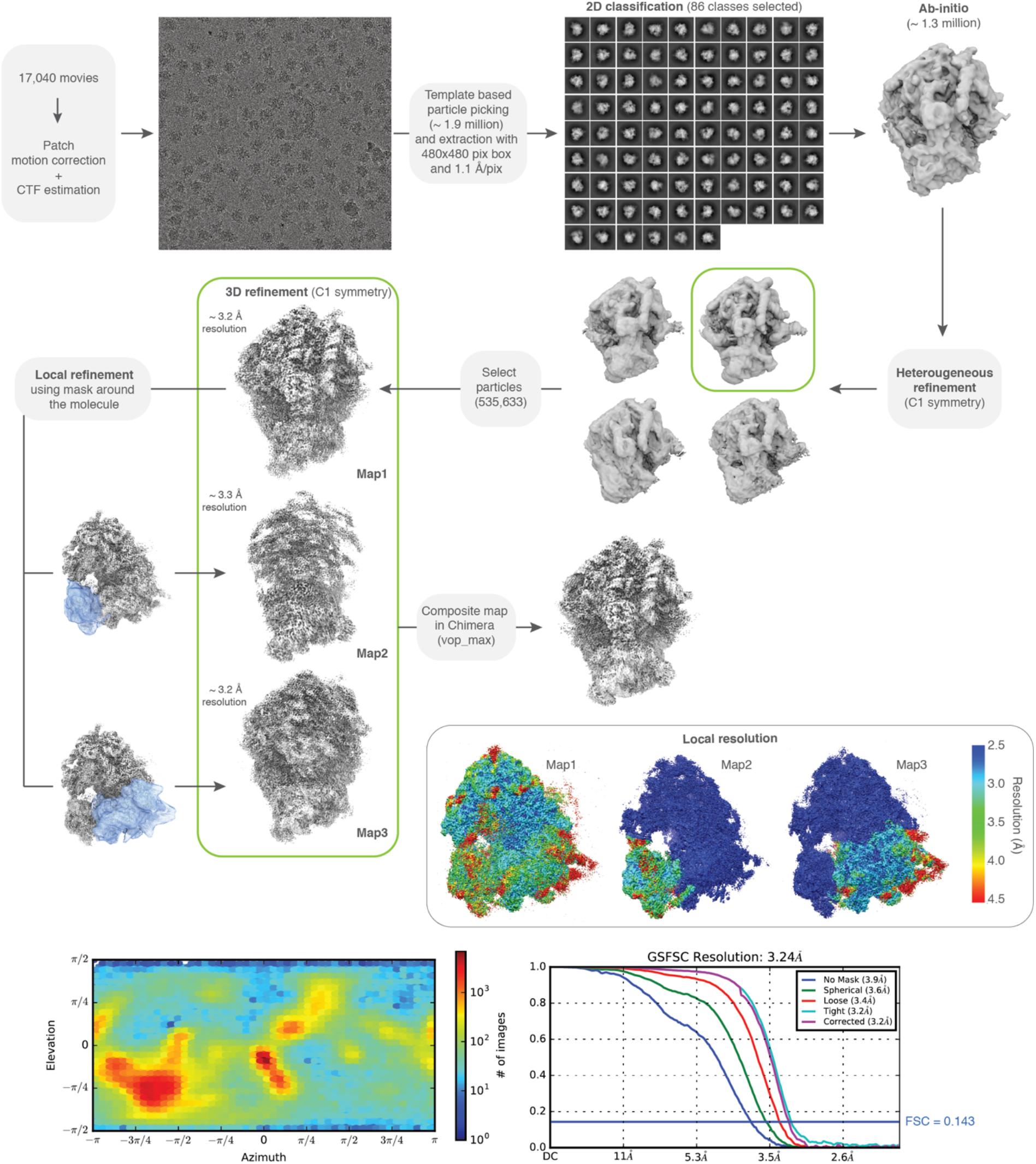
Processing pipeline of the 1 hpf zebrafish ribosome. All steps were done in Cryosparc v3.2.0. Maps are shown in gray, masks in blue. The orientation distribution plot for all particles contributing to Map1 and the Gold-Standard Fourier Shell Correlation (GSFSC) of the respective map is shown on the bottom. Local resolution maps were calculated for Map1, Map2 and Map3.

**Extended Data Fig. 3.**
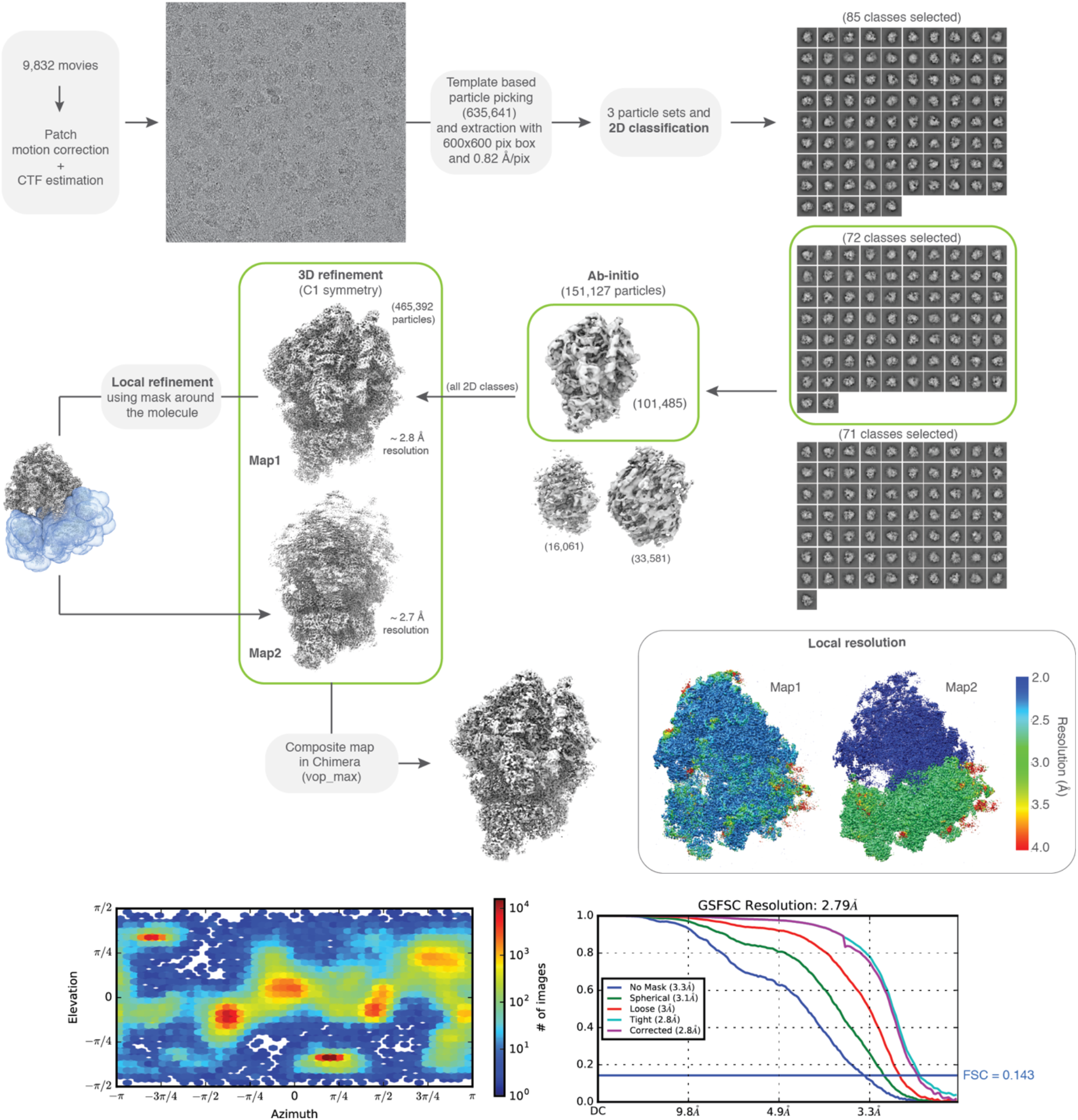
Processing pipeline of the *Xenopus egg* ribosome. All steps were done in Cryosparc v3.2.0. Maps are shown in gray, masks in blue. The orientation distribution plot for all particles contributing to Map1 and the Gold-Standard Fourier Shell Correlation (GSFSC) of the respective map is shown on the bottom right. Local resolution maps were calculated for Map1 and Map2.

**Extended Data Fig. 4.**
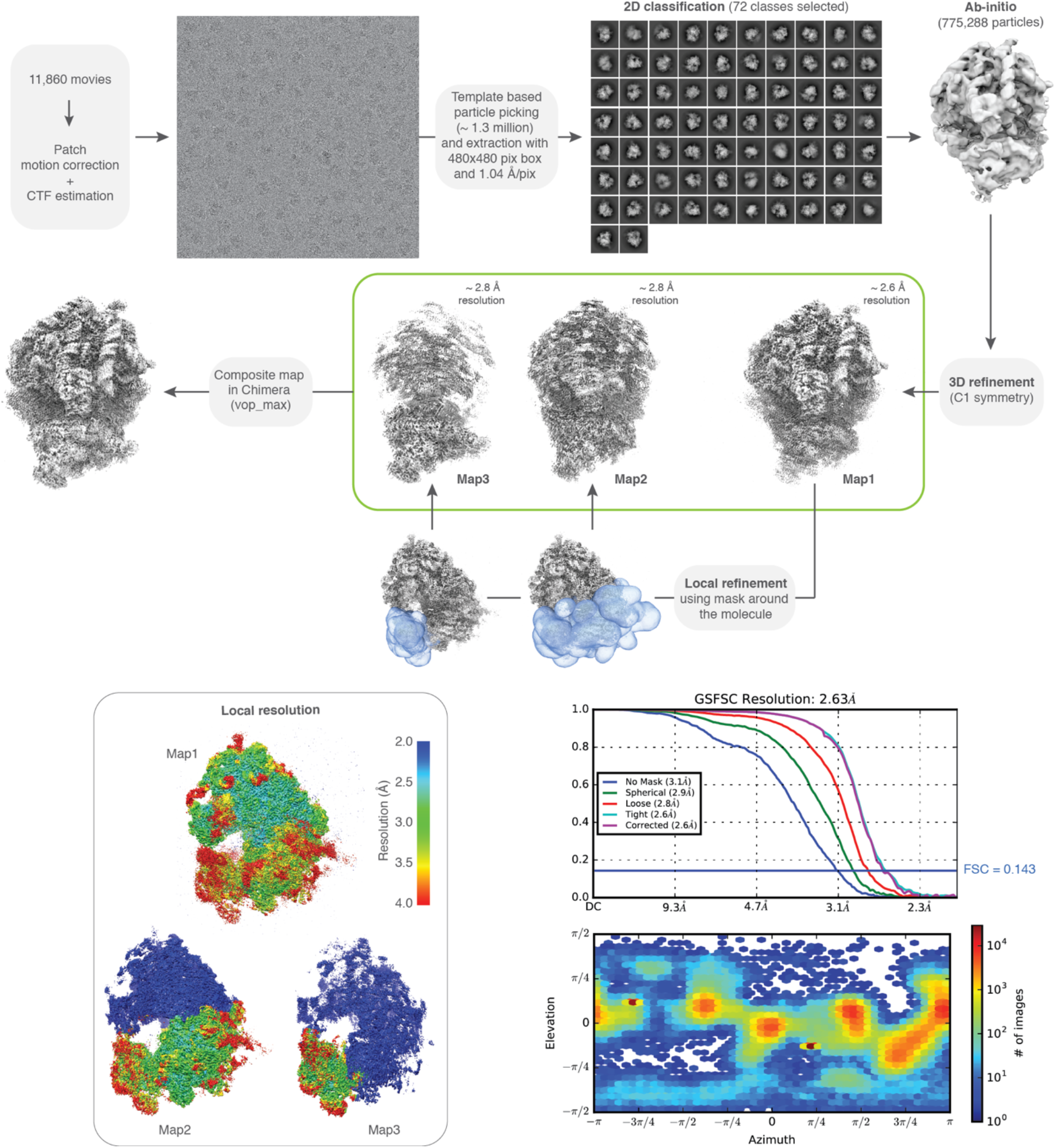
Processing pipeline of the 6 hpf zebrafish ribosome. All steps were done in Cryosparc v3.2.0. Maps are shown in gray, masks in blue. The orientation distribution plot for all particles contributing to Map1 and the Gold-Standard Fourier Shell Correlation (GSFSC) of the respective map is shown on the bottom-right. Local resolution maps were calculated for Map1, Map2 and Map3.

**Extended Data Fig. 5.**
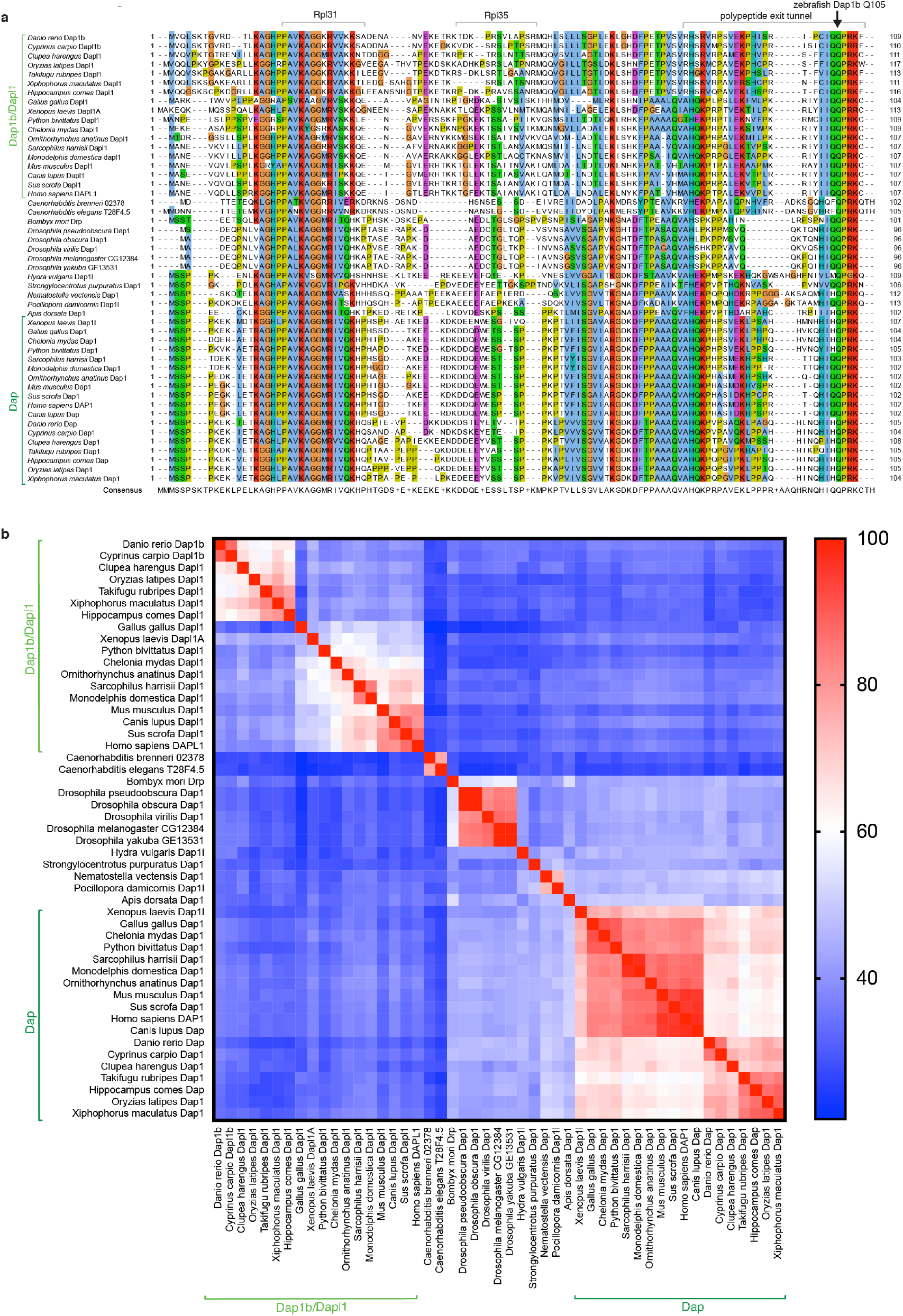
Analysis of amino acid sequence conservation of the Dap protein family. **a**, Protein sequence alignment illustrates sequence motifs shared between all Dap family proteins. Vertebrates have two paralogues, namely Dap1b/Dapl1 (light green) and Dap (dark green). Invertebrates only encode one homolog (Dap1) that clusters in between Dap1b/Dapl1 and Dap proteins. **b**, Percentage identity matrix illustrating the homology of Dap1b/Dapl1 (light green), Dap (dark green) and ancestral Dap1 proteins across different species. Red, 100% sequence identity; blue, 25% sequence identity.

**Extended Data Fig. 6.**
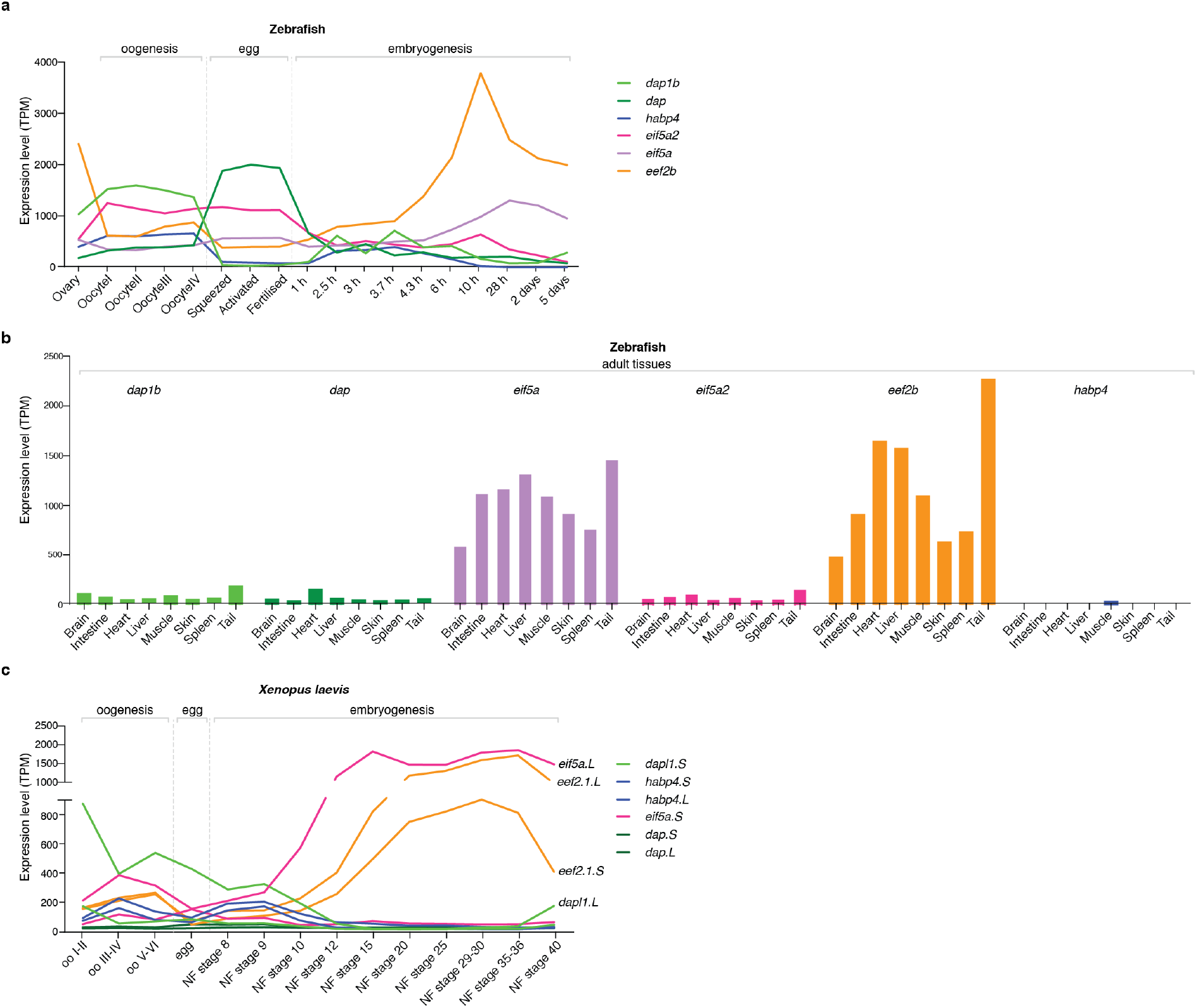
RNA expression of ribosome-associated factors during oogenesis, egg stages, embryo development and across adult tissues in zebrafish and *Xenopus*. (**A**) mRNA expression levels of zebrafish *eif5a/eif5a2* (purple), *eef2* (orange), *dap1b/dap* (green) and *habp4* (blue). PolyA+ RNA-seq during zebrafish oogenesis (oocyte stages I-IV), egg-stages (un-activated, activated, fertilized) and embryogenesis (2–4 cell, 256 cell, 1000 cell, oblong, dome, shield, bud, 1 day, 2 days, 5 days post-fertilization)^24,85^; TPM, transcripts per million. **b**, mRNA expression levels of ribosome-associated factors in zebrafish adult tissues. Amongst all the tissues tested, only general translation factors *eif5a* and *eef2b* transcripts are highly expressed in adult tissues. TPM, transcripts per million. **c**, mRNA expression levels of all paralogs of *Xenopus eif5a, eef2, dap, dapl1* and *habp4* derived from riboMinus-seq data^86^. TPM, transcripts per million.

**Extended Data Fig. 7.**
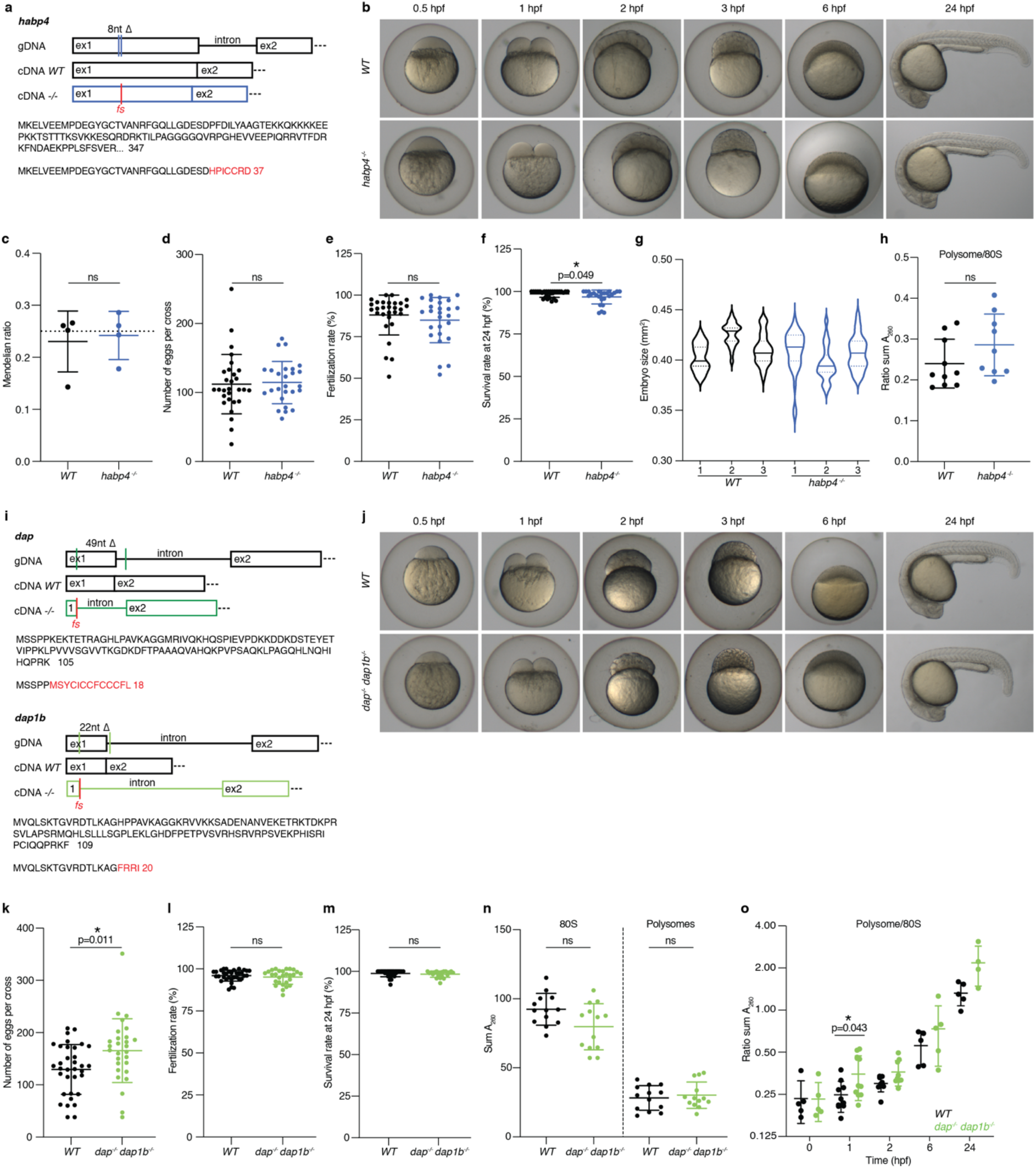
Characterization of *habp4*^*-/-*^ *and dap*^*-/-*^, *dap1b*^*-/-*^ zebrafish mutants. **a**, Scheme showing a deletion of 8 nucleotides in exon 1 of *habp4* which leads to a truncated protein of 37 amino acids. **b**, Early development of *habp4*^*-/-*^ embryos compared to wild type (*WT*). **c**, Mendelian ratio of fin-clips from a cross of *habp4*^+/-^ parents (25% of *habp4*^-/-^ and *WT* fish are expected (dotted line), n=4 crosses). **d**, Number of eggs laid per *WT* (n=29 crosses) or *habp4*^*-/-*^ (n=26 crosses) female. **e**, Fertilization rates of *WT* (n=29 crosses) and *habp4*^*-/-*^ (n=26 crosses) fish. **f**, Survival rate of *WT* (n=29 crosses) and *habp4*^*-/-*^ (n=25 crosses) 24 hpf embryos. **g**, Size (in square millimeters) of *WT* and *habp4*^*- /-*^ 6 hpf embryos obtained after setting up 3 different pairs of fish for mating (X axis). Significant differences in size (*p-value* <0.05, Kruskal-Wallis test) occur within (2/3 *WT*, 1/3 *habp4*^*-/-*^) and between genotypes (4/9 comparisons) (n = 31, 22 and 35 embryos for *WT* pairs 1-3, respectively; n=36, 24 and 33 embryos for *habp4*^*-/-*^ pairs 1-3, respectively). **h**, Quantification of polysome-to-monosome ratios of polysome gradients from *WT* and *habp4*^-/-^ 1 hpf embryos (n=9). **i**, Scheme showing deletions in exon 1 and intron 1 of *dap* (49 nucleotides) and *dap1b* (22 nucleotides) leading to truncated proteins of 18 (Dap) and 20 (Dap1b) amino acids. **j**, Early embryo development of *dap*^*-/ -*^, *dap1b*^-/-^ mutants compared to *WT*. **k**, Number of eggs laid by *WT* (n=34 crosses) or *dap*^*-/-*^, *dap1b*^*-/-*^ (n=28 crosses) fish. **l**, Fertilization rates of *WT* (n=34 crosses) and *dap*^*-/ -*^, *dap1b*^-/-^ (n=28 crosses) fish. **m**, Survival rate of *WT* (n=28 crosses) and *dap*^*-/ -*^, *dap1b*^-/-^ (n=21 crosses) 24 hpf embryos. **n**, Sum of A_260_ in polysome gradient analyses of monosomes and polysomes (n=13) of *WT* and *dap*^*-/-*^, *dap1b*^-/-^ 1 hpf embryos. **o**, Quantification of polysome-to-monosome ratios (in Log2 scale) at different stages of development of *WT* and *dap*^*-/-*^, *dap1b*^-/-^ eggs and embryos (0 h: n=5; 1 hpf: n=10; 2 hpf: n=8; 6 hpf: n=5; 24 hpf: n=4). Data in (c-f, h, k-o) are represented as scatter dot plots with means and SD. Data in g are represented as violin plot with median and quartiles. Significance was determined using Mann-Whitney test (c-f, h, k-o) or Kruskal-Wallis test (g). ns=not significant, * *p-value* < 0.05, hpf=hours post fertilization, fs=frameshift.

**Extended Data Fig. 8.**
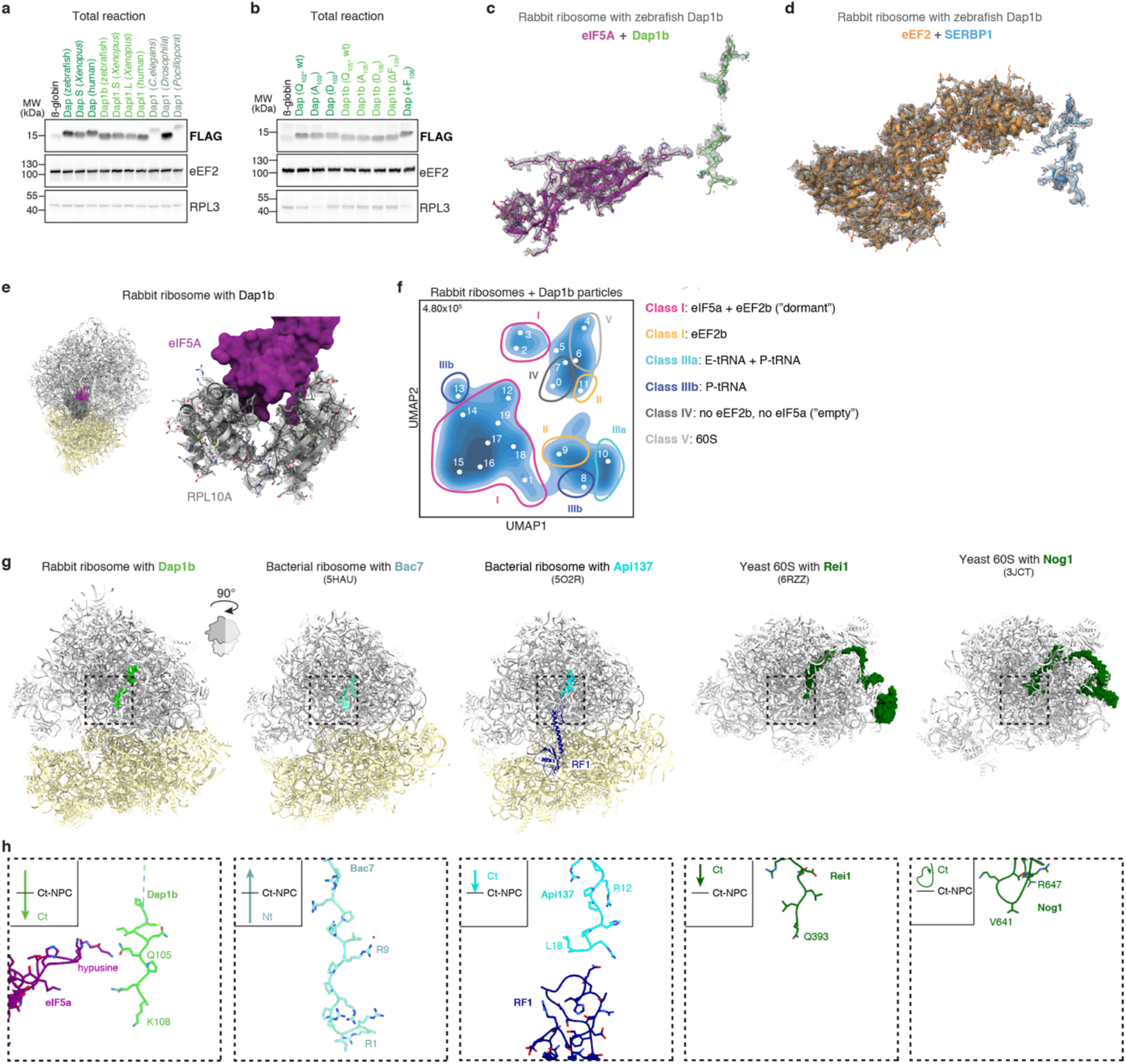
Recombinant Dap1b binds to the polypeptide exit tunnel of rabbit ribosomes. **a**, Western blot of the total *in vitro* translation reaction shown in Fig. 4D. **b**, Western blot of the total *in vitro* translation reaction shown in Fig. 4F. **c-d**, Densities of eIF5A and zebrafish Dap1b (c), and eEF2 and SERBP1 (d) modules in the rabbit ribosome with recombinant zebrafish Dap1b. **e**, The L1 stalk protein RPL10A (in gray) interacts with eIF5A (in dark magenta) in the rabbit ribosome with recombinant zebrafish Dap1b. An overview of the RPL10A position on the rabbit ribosome is shown on the left; RPL10A density is shown in mesh (right). **f**, Latent space representation of ribosomal particles of the dataset containing rabbit ribosomes with recombinant zebrafish Dap1b protein as a UMAP embedding after training a cryoDRGN latent variable model. Classes are depicted with circles in Roman numbers, map volumes are indicated with Arabic numbers. Total particle number is shown on the top left of the graph. **g**, Structures of ribosomes with proteins and peptides inserted into the polypeptide exit tunnel. From left to right: zebrafish Dap1b, Bac7 (5HAU^16^), Api137 (5O2R^57^), Rei1 (6RZZ^58^) and Nog1 (3JCT^61^). Models were superimposed using the command *mmaker* in ChimeraX and clipped to have a better view of the polypeptide exit tunnel. Boxed areas (dashed boxes) are shown at higher magnification in H. **h**, Detail of the peptidyl-transferase center (PTC) of the ribosomes shown in G. Schemes on the top-left of each box illustrate the relative position of the factor within the PTC in relation to the C-terminal residue of the nascent peptide chain (NPC). Nt refers to N-terminus, Ct corresponds to C-terminus.

**Extended Data Fig. 9.**
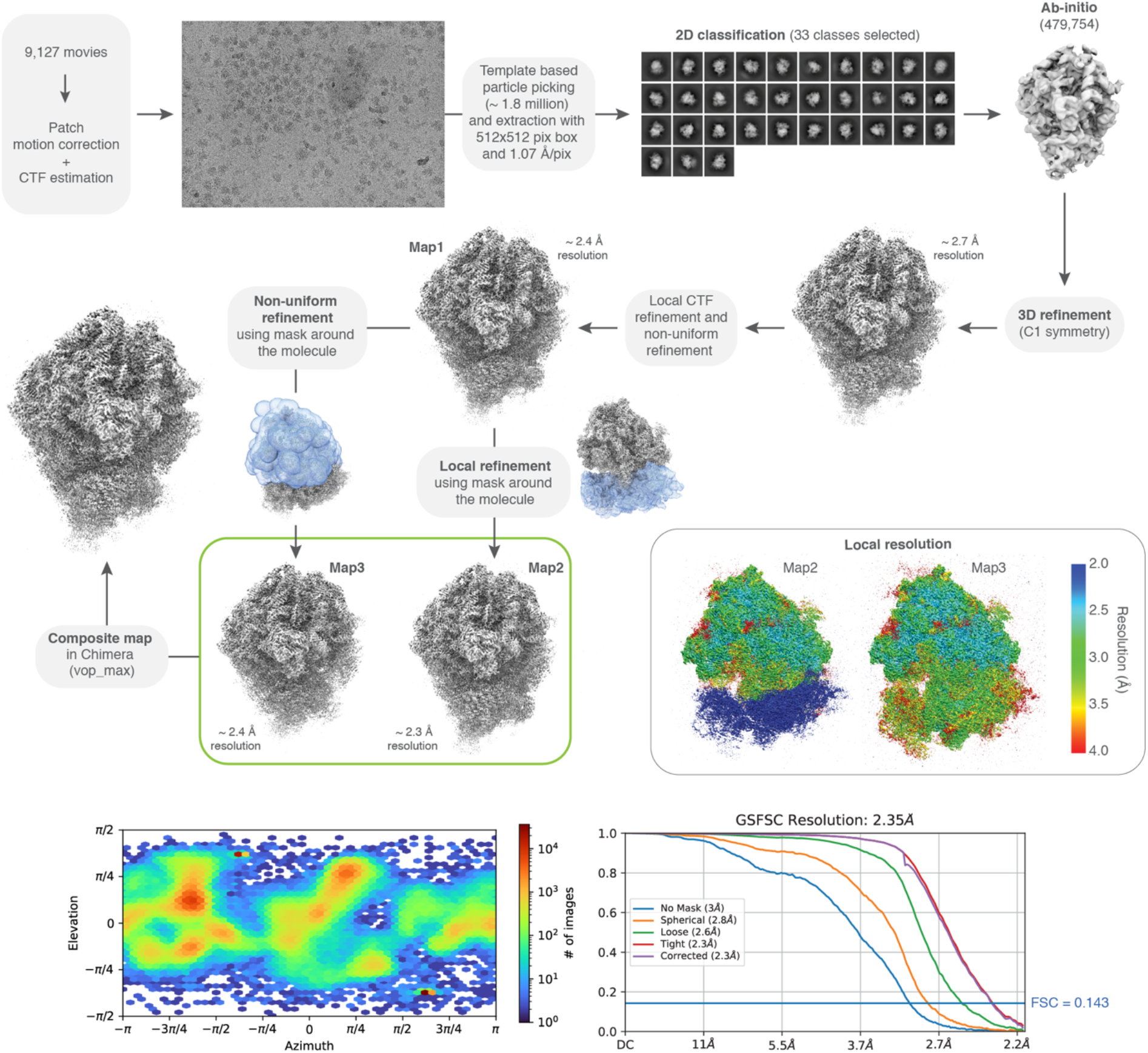
Processing pipeline of the rabbit ribosome with recombinant zebrafish Dap1b. Processing pipeline for obtaining an 80S density map of the rabbit ribosome with Dap1b. Local resolution maps were calculated for Map2 and Map3. The orientation distribution plot for all particles contributing to Map1 and the Gold-Standard Fourier Shell Correlation (GSFSC) of the respective map is shown on the bottom.

**Extended Data Table 1.** Cryo-EM data collection and refinement statistics of the ribosome structures presented in this study.

|  | 1 hpf zebrafish | 6 hpf zebrafish | *Xenopus* egg | Rabbit + Dap1b |
| --- | --- | --- | --- | --- |
| PDB ID | 7OYA | 7OYB | 7OYC | 7OYD |
| **Data collection** |  |  |  |  |
| Microscope | Thermo Fisher Titan Krios |  |  |  |
| Acceleration Voltage (kV) | 300 |  |  |  |
| Detector | Falcon 3 EC | Falcon 3 EC | Falcon 3 EC | K3 |
| Magnification (nominal) | 81,000 | 81,000 | 105,000 | 81,000 |
| Defocus range ($\mu\text{m}$) | 0.5-4 | 0.5-3 | 0.2-2 | 0.4-3.2 |
| Calibrates pixel size ($\text{\AA}/\text{px}$) | 1.1 | 1.04 | 0.82 | 1.07 |
| Electron exposure ($\text{e}/\text{\AA}^2$) | 43 | 48 | 40 | 50 |
| Exposure rate ($\text{e}/\text{\AA}^2/\text{s}$) | 43 | 48 | 40 | 50 |
| Number of frames per movie | 39 | 39 | 39 | 41 |
| Collection software | Cryosparc 3.2.0 | Cryosparc 3.2.0 | Cryosparc 3.2.0 | Cryosparc 3.2.0 |
| Number of micrographs | 17,040 | 11,860 | 9,832 | 9,127 |
| Initial particle number | 1,961,364 | 1,285,670 | 635,641 | 1,808,120 |
| Final particle number | 535,633 | 775,288 | 465,392 | 479,754 |
| Map resolution ($\text{\AA}$, FSC=0.143) | 3.2 | 2.6 | 2.8 | 2.4 |
| **Refinement** |  |  |  |  |
| Software | Phenix 1.17.1 | Phenix 1.17.1 | Phenix 1.17.1 | Phenix 1.17.1 |
| Initial model(s) | 4UG0<br>6MTE<br>5DAT | 4UG0 | 4UG0<br>6MTE<br>5DAT | 6MTE<br>5DAT<br>5GAK |
| Correlation coefficient ($\text{CC}_{\text{mask}}$) | 0.83 | 0.84 | 0.82 | 0.76 |
| Map sharpening factor ($\text{\AA}^2$) | -108.5 | -89.2 | -73.4 | -50.2 |
| Model composition (chains) | 82 | 77 | 81 | 85 |
| Non-hydrogen atoms | 199,914 | 194,870 | 198,952 | 213,014 |
| Protein residues | 11,179 | 10,795 | 11,387 | 12,293 |
| nucleotides | 5,130 | 5,035 | 5,006 | 5,316 |
| Ligands | MG: 202<br>ZN: 7 | MG: 203<br>ZN: 7 | MG: 134<br>ZN: 7 | ZNA: 1<br>MG: 201<br>B8N: 1<br>ZNK: 1<br>ZN: 5<br>4AC: 2 |
| **B factors ($\text{\AA}^2$, min/max/mean)** |  |  |  |  |
| Protein | 4.4 / 281.9 / 53.3 | 12.8 / 350.6 / 51.7 | 4.4 / 300.2 / 53.4 | 4.4 / 244.8 / 82.7 |
| RNA | 13.9 / 234.1 / 53.1 | 13.9 / 205.7 / 56.6 | 13.9 / 205.7 / 54.6 | 20.0 / 333.2 / 81.7 |
| Ligands | 7.8 / 102.4 / 28.4 | 7.8 / 102.4 / 27.0 | 7.8 / 102.4 / 25.1 | 25.8 / 120.5 / 57.1 |
| **RMS deviations** |  |  |  |  |
| Bond length ($\text{\AA}$) (#>4 sigma) | 0.003 | 0.003 | 0.003 | 0.005 |
| Bond angle ($^\circ$) (#>4 sigma) | 0.620 | 0.614 | 0.619 | 0.714 |
| **Validation** |  |  |  |  |
| MolProbity score | 1.59 | 1.72 | 1.54 | 1.68 |
| Clashscore | 6.04 | 8.01 | 6.95 | 6.96 |
| Poor Rotamers (%) | 1.30 | 1.32 | 0.76 | 1.06 |
| CB deviations (%) | 0.01 | 0.00 | 0.00 | 0.01 |
| CABLAM outliers (%) | 2.35 | 2.36 | 2.27 | 2.25 |
| Favored (%) | 97.03 | 96.86 | 97.10 | 96 |
| Allowed (%) | 2.97 | 3.14 | 2.90 | 4 |
| Disallowed (%) | 0 | 0 | 0 | 0 |

**Extended Data Table 2.**
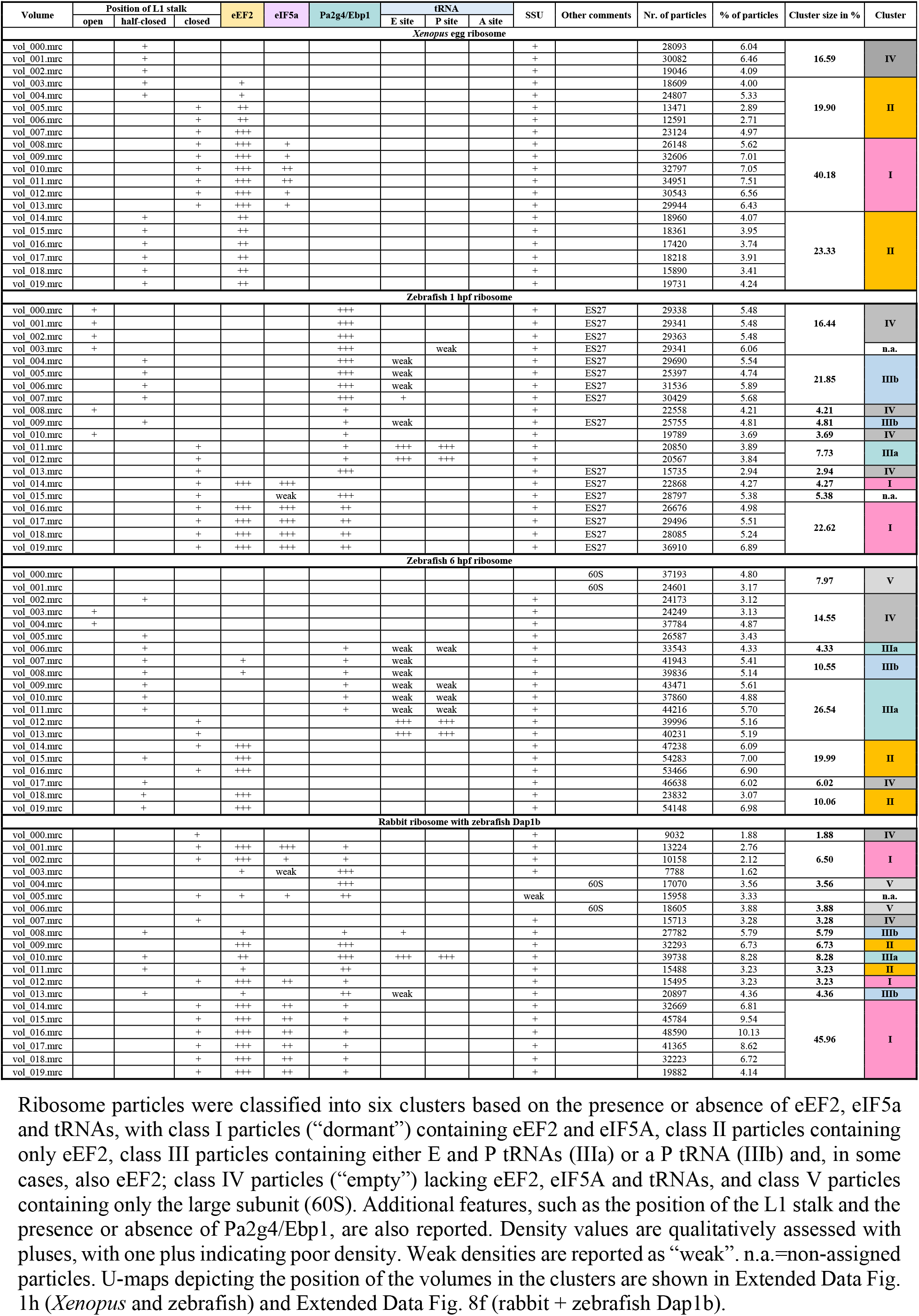
Classification of the map volumes obtained with CryoDRGN.

**Extended Data Table 3.**
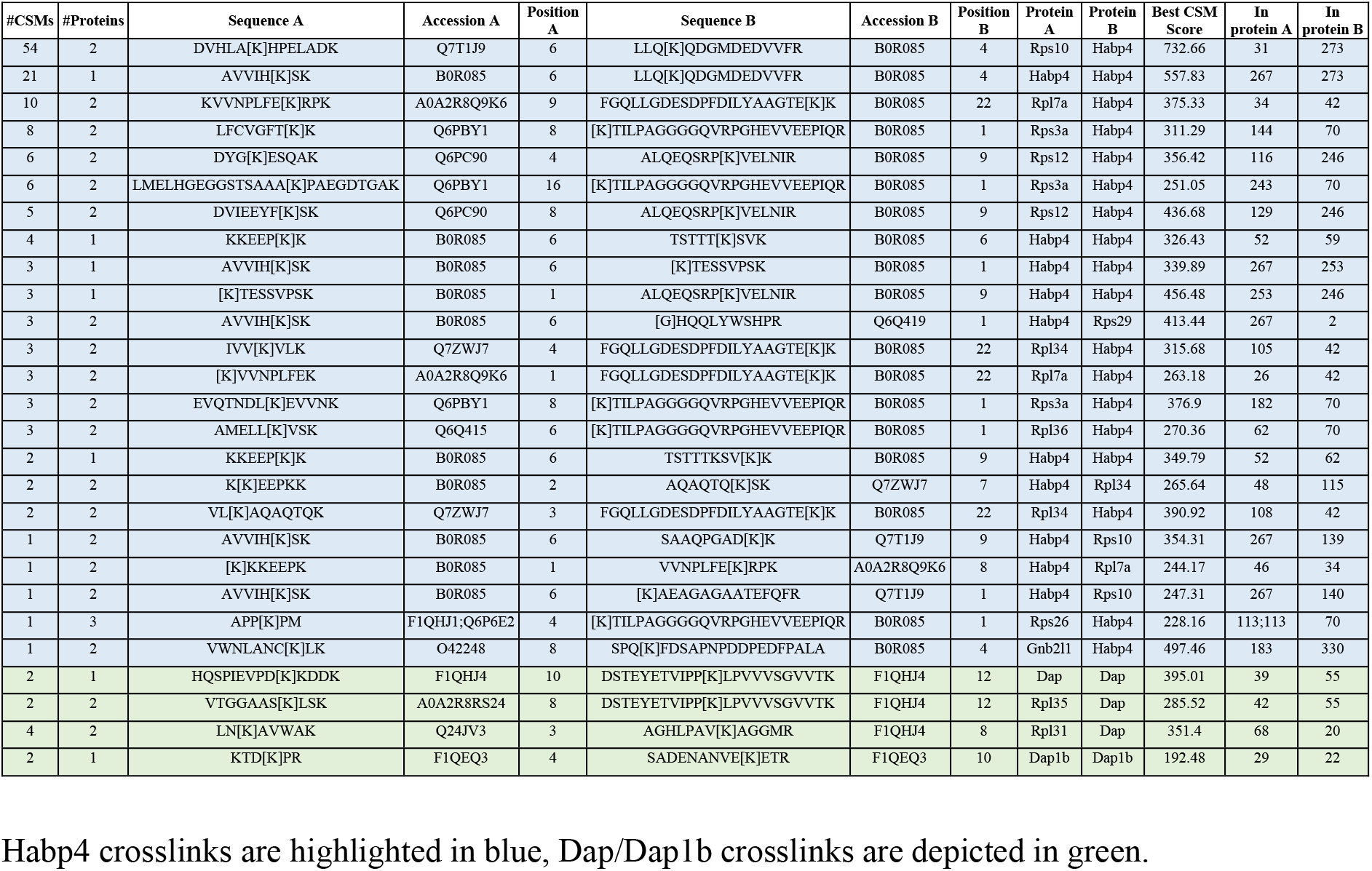
Crosslinks identified in ribosome samples from 1 hpf zebrafish embryos.

**Extended Data Table 4.**
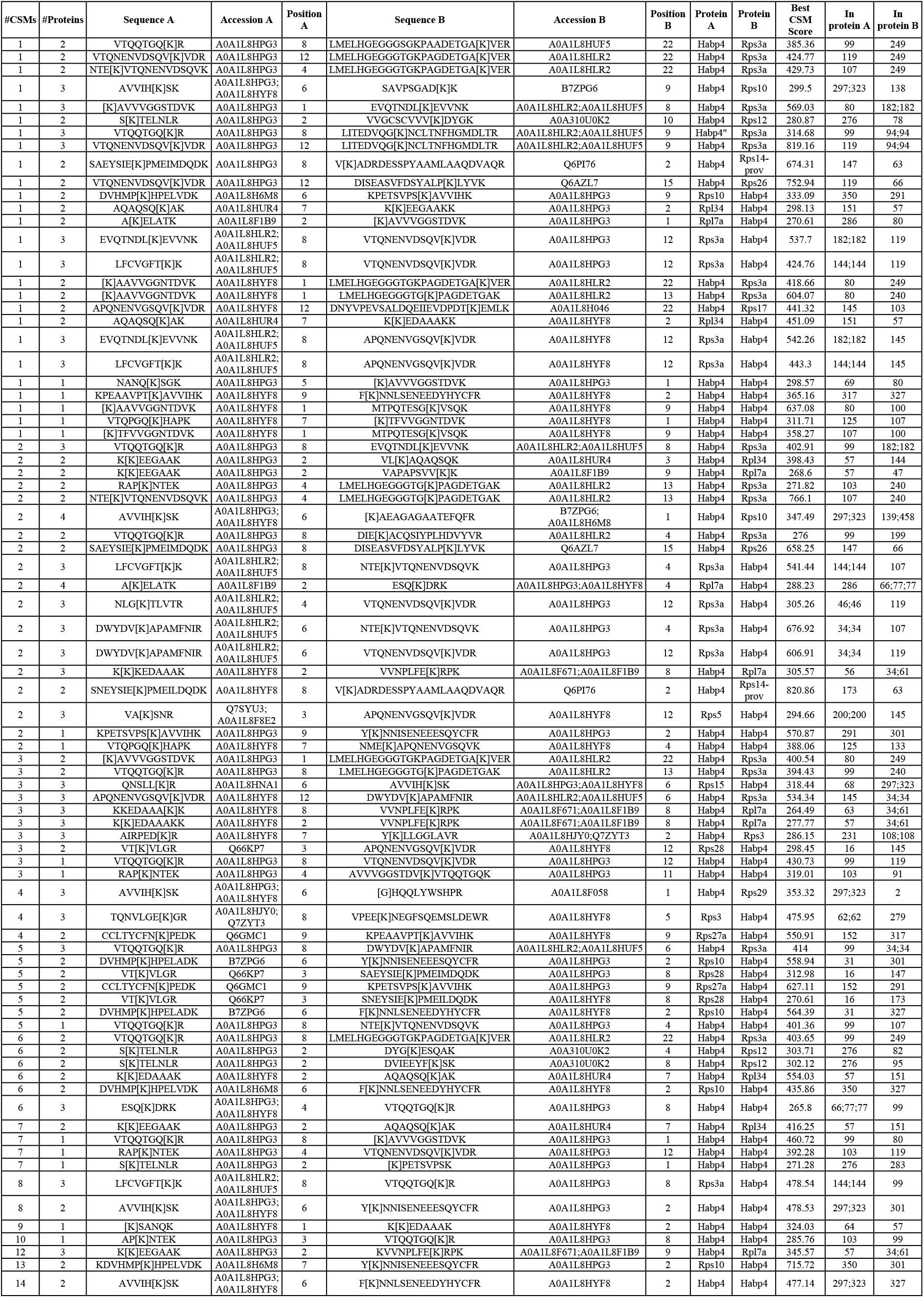
Crosslinks identified for Habp4 in ribosome samples from *Xenopus* eggs.

**Extended Data Table 5.**
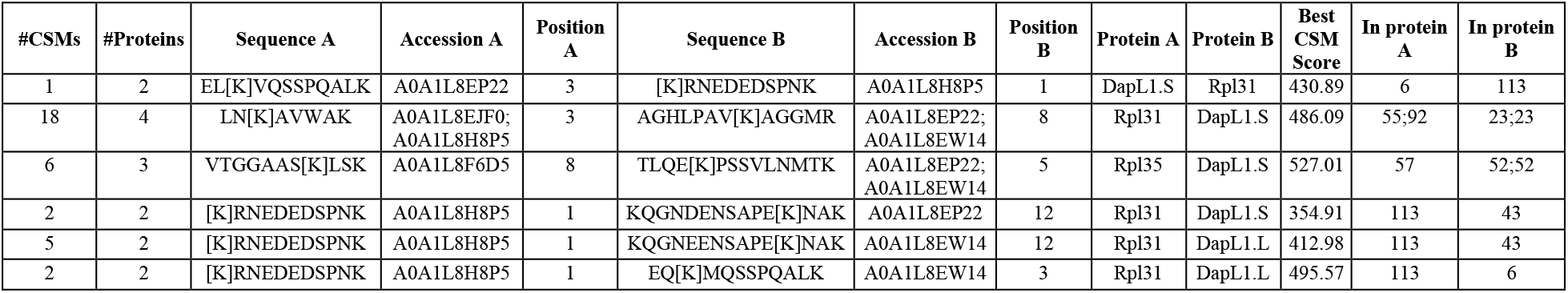
Crosslinks identified for Dapl1 in ribosome samples from *Xenopus* eggs.

**Extended Data Table 6.**
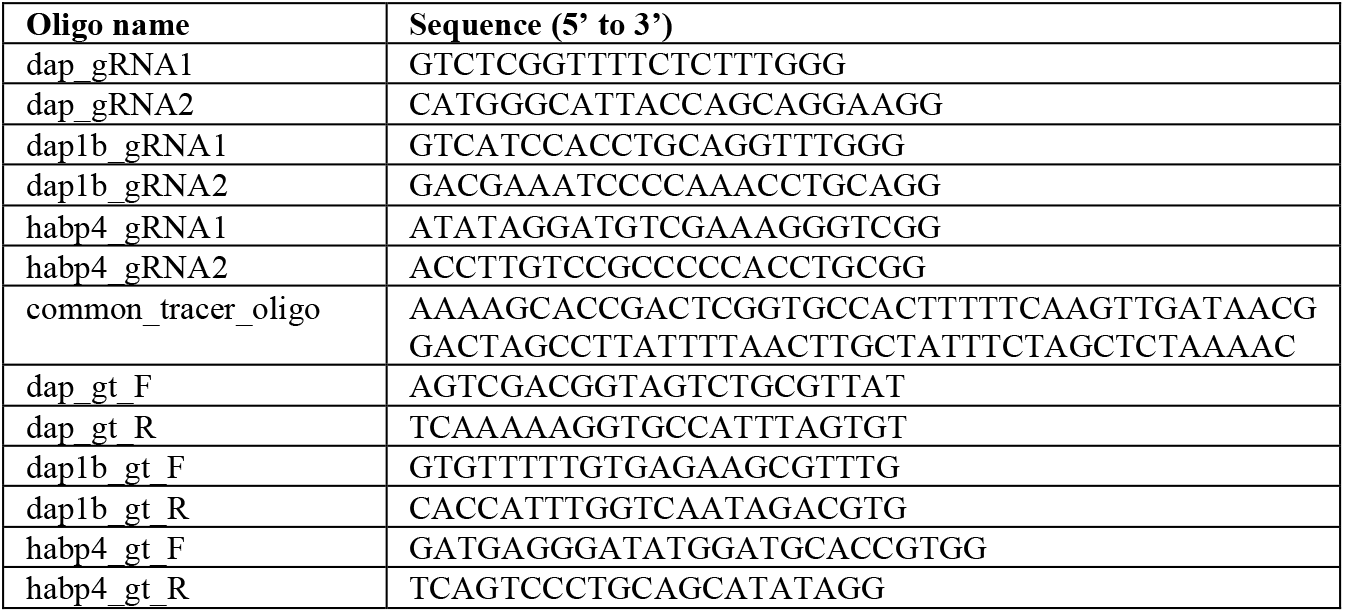
Primers used in this study.

## Notes

### Competing Interest Statement

The authors have declared no competing interest.

